# Reciprocal control of the stomach and intestine by viscerosensory neurons in caudal brainstem

**DOI:** 10.64898/2026.09.21.753299

**Authors:** Naz Dundar, Ilayda Alkislar, Brooke C. Jarvie, Ho Namkung, Kathryn Xie, Mahekdeep Kaur, Queenie Li, Anagh S. Ravi, Alejandro López-Cruz, Antoinette Spina, Longhui Qiu, Truong Ly, Jun Y. Oh, Kevin Yackle, Zachary A. Knight

## Abstract

Digestion requires precisely timed transit of food through the gastrointestinal tract. Here we show that emptying of food from the stomach occurs in discrete pulses, separated by pauses that allow the proximal duodenum to clear before the next bolus arrives. By combining X-ray fluoroscopy of the gut with calcium imaging of the brainstem, we identify a population of neurons defined by expression of neuropeptide FF (NPFF) that tracks these duodenal dynamics. Stimulation of NPFF neurons induces a coordinated response that inhibits gastric emptying and accelerates intestinal transit, thereby clearing the proximal intestine for the next bolus of food. These neurons also bidirectionally control food intake through cholinergic signaling in the gut. These findings reveal a mechanism by which brainstem transforms the sensory detection of nutrients in the duodenum into the opposing control of the stomach and intestine. They also establish an experimental approach for dissecting gut-brain signaling in behaving animals.

## Introduction

The gastrointestinal (GI) tract is highly dynamic^1–8^, with coordinated motion of the stomach and intestine occurring on timescales from seconds to minutes^4,9–13^. This coordination is critical for the digestion and absorption of food, and its dysregulation is associated with a variety of GI disorders^1,3,7,14,15^. Previous work has shown that GI motility is controlled by the interaction between intrinsic circuitry in the gut, which generates rhythmic contractions^11,16–22^, and top-down modulation by the brainstem^23–29^, which is relayed by the vagus nerve. This top-down modulation is poised to organize GI rhythms across organs^30^, but little is known about how this multi-organ coordination is achieved during natural ingestion.

Addressing this question requires defining how the movement of food through the GI tract is represented in the brain and used to control GI function in an awake, behaving animal. While conceptually straightforward, this has been challenging to do in practice for two reasons. The first is that the key neurons that directly receive information from the gut are located in the caudal nucleus of the solitary tract (cNTS) in the brainstem^24,31–34^, and recordings of cNTS neurons in awake animals have historically not been possible. The second is that commonly-used methods for measuring GI motility are based on either endpoint assays^35–37^, which lack spatial and temporal resolution, or ex vivo preparations^38–40^, which cannot resolve the natural flow of ingested food through the gut.

We reasoned that we could close this gap by combining two advances: the ability to monitor the activity of viscerosensory neurons in the caudal brainstem of awake mice, which we recently demonstrated^41,42^, and X-ray fluoroscopy, which allows GI motility to be visualized in real time and during behavior^43–45^. Here, we use this strategy to reveal how the brainstem monitors GI dynamics and then transforms them into the reciprocal control of the stomach and intestine, enabling the precisely timed transit of food.

## Results

### Pulsatile dynamics of food transit through the gut can be visualized by X-ray fluoroscopy

We first set out to characterize the dynamics of the transit of food through the GI tract in awake mice. To do this, we used X-ray fluoroscopy, which enables the visualization of GI dynamics by generating X-ray movies of a contrast agent as it passes through the gut^43,45^. Mice were equipped with intragastric (IG) catheters, head-fixed in the imaging setup, and then received infusion of a barium-nutrient mixture (0.5 mL over 5 min, 30% barium in either 10% intralipid or 24% glucose, Fig. 1A, B, Fig. S1B). We then monitored this mixture as it transited through the gut.

**Figure 1.**
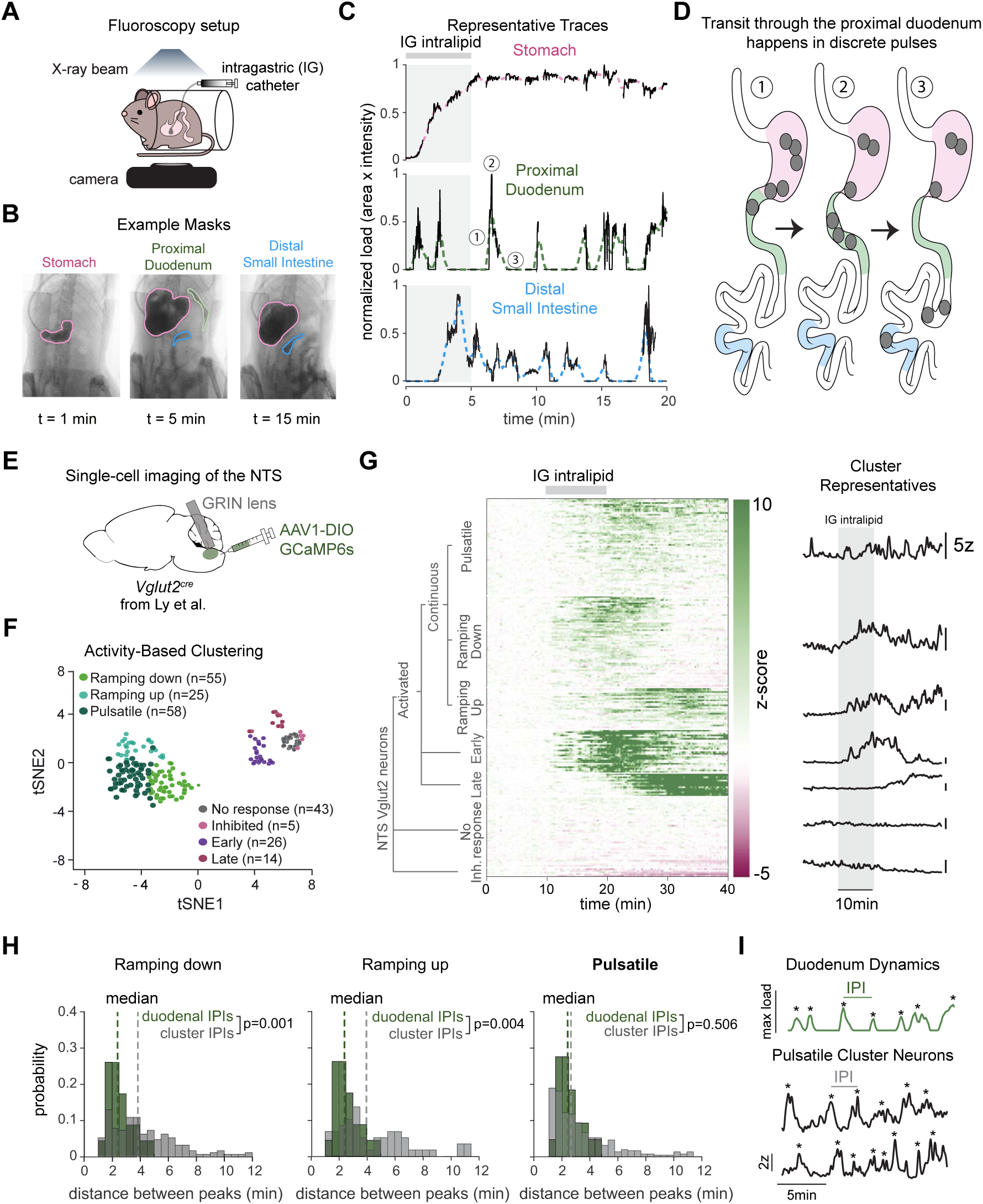
A subpopulation of NTS neurons displays dynamics that mirror duodenal motility. **A.** Schematic of X-ray fluoroscopy setup. Mice were infused intragastrically (IG) with a barium/intralipid mixture. **B.** Representative fluoroscopy images at t = 1, 5 and 15 min following the start of infusion with stomach (pink), proximal duodenum (green) and distal small intestine (blue) masks outlined. The distal small intestine mask was placed at varying positions across animals. **C.** Representative normalized load traces for the stomach, proximal duodenum and distal small intestine from a single mouse during IG infusion (gray bar). Colored dashed lines show smoothed, interpolated traces. Gray shading indicates the IG infusion period. Numbers (1–3) indicate phases of a pulse of duodenal filling, corresponding to the schematic in D. **D.** Schematic illustrating that contents transit into the proximal duodenum in discrete pulses separated by pauses, rather than continuous flow. Numbers correspond to the phases of a pulse indicated in C. **E.** Single-cell calcium imaging of NTS glutamatergic neurons (Vglut2^cre^; AAV1-DIO-GCaMP6s, n = 5 mice, 226 neurons) reanalyzed from Ly et al^41^. **F.** tSNE visualization of all recorded neurons colored by cluster identity. **G.** Activity-based clustering (see Fig. S2) of NTS Vglut2 neurons during IG infusion. Heatmap shows z-scored activity for all neurons sorted by cluster. Right, representative trace for each cluster. Scale bars, 5z, 10 min. **H.** Interpeak interval (IPI) distributions for duodenal traces (green) and neural clusters (gray); Ramping down, Ramping up, and Pulsatile. Lines indicate medians for duodenal IPIs (green) and neural cluster IPIs (gray). p values indicate comparison of duodenal and cluster IPI distributions. **I.** Top, Representative duodenal fluoroscopy dynamics trace. Bottom, z-scored traces from individual Pulsatile cluster neurons. Asterisks indicate peaks. Scale bars, maximum load (top), 2z (bottom), 5 min.

We observed an increase in gastric contrast as the stomach filled during the infusion, which then slowly decayed over tens of minutes after the infusion was complete (Fig. 1C, Fig. S1C, D). In the proximal duodenum, contrast appeared with a latency of 48.2 ± 14.4 s relative to stomach filling, indicating that gastric emptying begins rapidly during the IG infusion. This contrast then appeared in the more distal small intestine after a minute or more (latency 138.8 ± 43.3 s, Fig. S1E), confirming that X-ray fluoroscopy can be used to track the transit of food.

We noticed that filling of the proximal duodenum occurred in discrete pulses (Fig. 1C, D), which were typically cleared before the next bolus was delivered from the stomach (interpeak interval (IPI): median 2.3 min, IQR 1.8–3.1 min, n = 5 mice). These pulsatile dynamics were marked by rapid filling of the proximal duodenum, followed by segmentation^46^ (i.e. local mixing), and then emptying to the more distal small intestine in small boluses (Video S1). Of note, these pulses were not initiated by changes in the rate of antral contractions, because these did not differ in rate significantly between emptying and non-emptying epochs (Fig. S1F, Video S2). Rather, these dynamics suggest tight coordination of gastric emptying and duodenal motility, possibly involving top-down control by the brain. We therefore set out to identify neurons in the brainstem that might be involved in this regulation.

### A subpopulation of NTS neurons exhibits pulsatile activation resembling GI dynamics

Information about gut motility is sensed by the vagus nerve and relayed to the caudal nucleus of the solitary tract (NTS)^31–33,47–49^. We therefore reasoned that there should be some cNTS neurons whose activity mirrors the pulsatile dynamics of duodenal motility that we observed by fluoroscopy. To investigate this, we reanalyzed data from a recent study that performed single-cell imaging of cNTS glutamatergic neurons (n = 226 neurons, 5 mice) in response to IG infusion of Intralipid in awake, head-fixed mice^41^, a protocol that mirrors our fluoroscopy setup (Fig. 1E-G). Hierarchical activity-based clustering (Fig. S2) revealed that most cNTS neurons were activated by IG infusion (n = 178), with a small number of non-responsive (n = 43) and very few inhibited cells (n = 5). We performed further clustering to subdivide the activated neurons into a major population that was active throughout the experiment (continuous, n = 138) and much smaller subgroups that showed a strong response either early (n = 26) or late (n = 14).

We reasoned that neurons tracking the filling of the proximal duodenum would (1) exhibit phasic responses corresponding to each cycle of gastric emptying and (2) show minimal ramping activity during the session, since GI contents do not accumulate over time (i.e., emptying typically occurs before the next bolus). We defined metrics that quantify these two requirements (“peakiness” and “baseline trend”) and then used these as the basis for a new round of clustering of the neurons active throughout the experiment, which yielded three subgroups of cells. One of these clusters, which we termed “pulsatile” (n = 58 neurons), showed calcium dynamics that closely matched the motility dynamics of the proximal duodenum measured by fluoroscopy (IPI median: 2.7 min vs 2.3 min; calcium vs. motility, KS test, p = 0.5), whereas the other two neural subsets had significantly different IPI distributions (median: 3.8 min, p = 0.004 and 3.9 min, p = 0.001, ramping up and down, respectively, Fig. 1H). This indicates that approximately 25% of cNTS neurons exhibit dynamic responses to IG infusion that could track intestinal motility (Fig. 1I).

### NPFF neurons track the entry of food into the duodenum

We next sought to identify a genetic marker that labels these pulsatile cNTS neurons. Previously, we reported the dynamics of two cNTS cell types, PRLH and GCG neurons, neither of which showed the pulsatile responses to IG infusion of Intralipid described above^42^. However, these two cell types are located more laterally in the cNTS, which is enriched for neurons that respond to oral and gastric stimulation^31^. In contrast, intestine-responsive cells are located more medially, and the vagal afferents that innervate the proximal intestine terminate most densely in a small region on the border between the cNTS and the area postrema known as the subpostrema^32,48^ (Fig. 2A). To identify a genetic marker for cells in the subpostrema, we analyzed published MERFISH data^50^ and ranked cNTS cell types by their proximity to this region, which revealed a glutamatergic cluster defined by expression of neuropeptide FF as the top hit (NPFF, Fig. S3). We generated an Npff-2A-Cre knockin mouse line to gain genetic access to these cells (Fig. S4); showed that Npff-Cre expression was restricted to the subpostrema in the caudal brainstem (Fig. S4A); and confirmed by RNAscope that Cre recombination faithfully recapitulates endogenous *Npff* expression (Fig. S4B).

**Figure 2.**
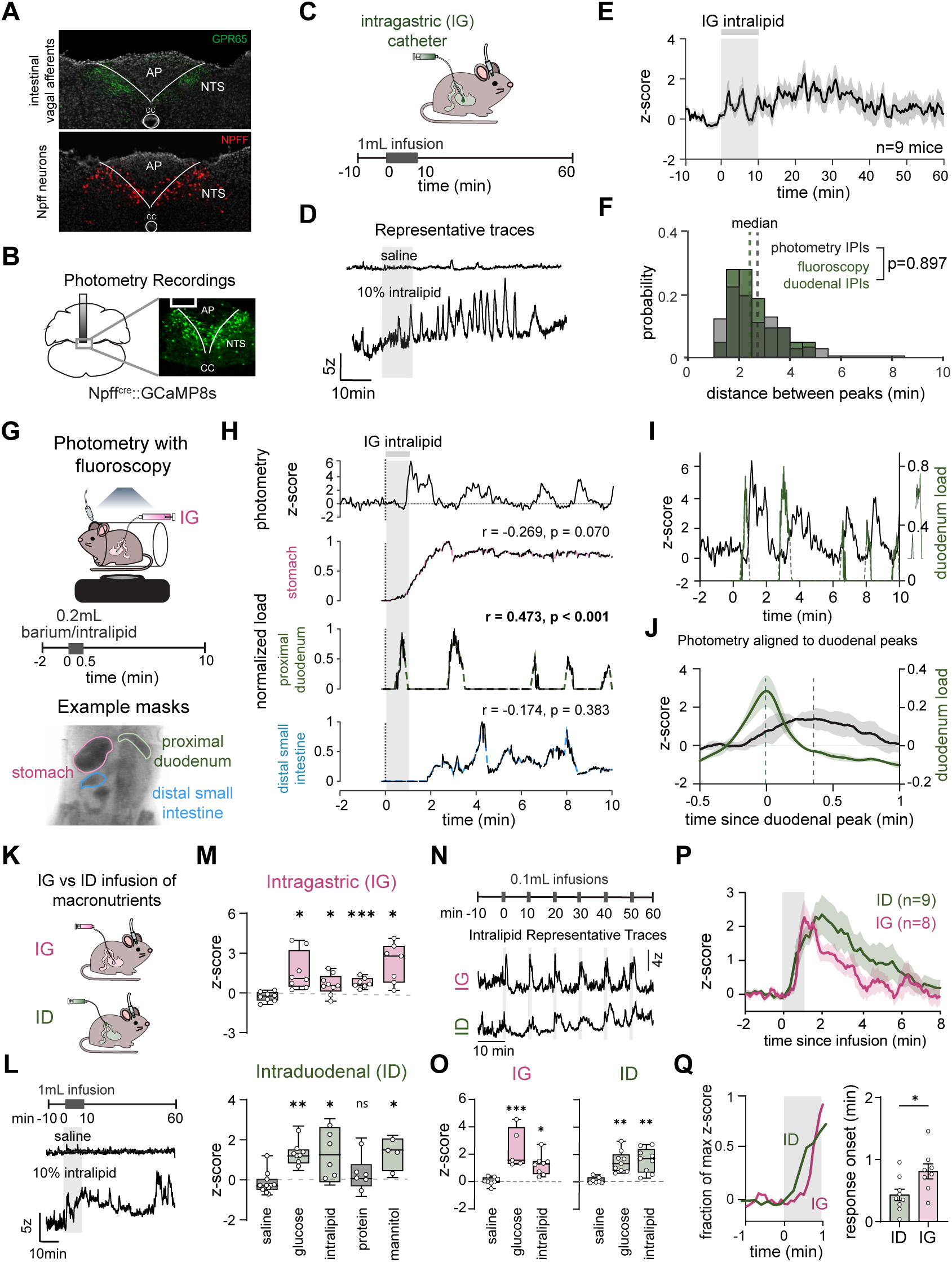
Subpostrema NPFF neurons respond to the filling of the proximal duodenum. **A.** Intestinal vagal afferent terminals and NPFF neurons are localized to the subpostrema. Top, vagal afferents labeled by *GPR65^Cre^::GCaMP6s* mice (green). (Bottom) NPFF neurons labeled by *Npff^cre^::tdTomato mice* (red). AP, area postrema. CC, central canal. **B.** *Npff^Cre^::GCaMP8s* mice were implanted with an angled optical fiber. Right, GCaMP8s expression. **C.** Intragastric (IG) infusion experiments. Mice were infused with 1mL of liquid over 10 min. **D.** Representative photometry traces from a single mouse during IG infusion of saline or 10% intralipid. Gray shading indicates infusion period. Scale bar, 5z, 10 min. **E.** Mean z-scored NPFF photometry activity during IG infusion of 10% intralipid (n = 9 mice). Gray shading indicates infusion period. **F.** Distribution of inter-peak intervals (IPIs) from NPFF photometry peaks during IG infusion (gray) compared to duodenal load IPIs from fluoroscopy recordings (green; from Fig. 1H). Dashed lines indicate medians. **G.** Simultaneous fiber photometry and X-ray fluoroscopy setup. Mice were infused IG with a 0.2mL 10% Intralipid/30% barium mixture. Bottom, example fluoroscopy masks for the stomach (pink), proximal duodenum (green), and distal small intestine (blue). **H.** Example simultaneous recording from a single mouse showing z-scored NPFF activity (top) and normalized load traces for the stomach, proximal duodenum, and distal small intestine (see Fig S5 for more examples). Pearson correlation coefficients between NPFF activity and organ load were calculated at a 22s lag (distance between NPFF and duodenal peaks, 2J) and compared to 1,000 iterations of circular shuffles. **I.** Example simultaneous trace from a single mouse (same as 2H) showing z-scored NPFF activity (black) and normalized duodenal load (green) aligned on the same time axis. **J.** Mean z-scored NPFF activity (black) aligned to duodenal load peaks (green) across all mice (n = 3). Dashed lines indicate duodenal (green) and NPFF peaks (black). **K.** Comparison of IG and intraduodenal (ID) infusion setups. **L.** Representative photometry traces from a single mouse during ID infusion of saline or 10% intralipid. Gray shading indicates infusion period. Scale bar, 5z, 10 min. **M.** Mean z-scored NPFF activity (0-10 min) during IG (top) and ID (bottom) infusions. IG: saline (n = 8), glucose (n = 8), Intralipid (n = 9), protein (n = 7), mannitol (n = 7). ID: saline (n = 5), glucose (n = 8), Intralipid (n = 6), protein (n = 6), mannitol (n = 4). **N.** Top, timeline of small bolus infusion experiment. Animals were infused with 0.1mL boluses of 10% intralipid every 10 minutes for an hour (6 infusions) IG (top) or ID (bottom). Bottom, representative photometry traces from a single mouse. Scale bar, 4z, 10 min. **O.** Mean z-scored NPFF neuron response per bolus during IG (left) and ID (right) infusions. Responses were averaged across all boluses (0-2 min) within each animal. IG: saline (n = 8), glucose (n = 5), Intralipid (n = 8). ID: saline (n = 7), glucose (n = 9), Intralipid (n = 9). **P.** Mean z-scored NPFF activity aligned to the start of each 0.1 mL bolus infusion for IG (green) and ID (pink). **Q.** Left, fraction of maximum response over time following infusion for IG and ID. Right, response onset (time to 2 s.d. above baseline) for IG vs ID infusions. Each dot represents a single mouse. *p < 0.05, **p < 0.01, ***p < 0.001, ns, not significant. Data are mean ± s.e.m. unless specified otherwise.

To determine whether these neurons respond to GI stimuli, we targeted GCaMP8s to NPFF neurons, implanted an optical fiber for calcium recordings by fiber photometry, and equipped mice with IG catheters (Fig. 2B, C). We found that IG infusion of Intralipid triggered pulsatile bursts in NPFF neuron activity throughout the recording session that closely resembled duodenal dynamics (Fig. 2D, E). For example, the distribution of inter-peak intervals for these transient peaks in calcium activity was indistinguishable from the dynamics of duodenal emptying measured in a separate experiment by fluoroscopy (IPI median 2.4 vs 2.3 min; calcium vs. motility, KS test, p = 0.9, Fig. 2F), suggesting that these peaks may correspond to individual duodenal filling events.

To test this hypothesis, we next combined fiber photometry and fluoroscopy so that we could monitor NPFF neuron activity and GI dynamics simultaneously in the same animal (Fig. 2G). This revealed that individual duodenal filling events were consistently followed by transient increases in NPFF neuron activity (Fig. 2H-J, Fig. S5, Video S3). These peaks in duodenal fill and calcium dynamics were consistently offset by 22.8 ± 4.1 s, and, after correcting for this lag, NPFF activity was highly correlated with duodenal load, which we confirmed by comparison to circularly shuffled controls (NPFF activity vs duodenal load, r = 0.536 ± 0.038, all p < 0.001). In contrast, NPFF activity either showed no correlation or, in some animals, a negative correlation with stomach and distal small intestine load. This negative correlation likely reflects the fact that proximal duodenum fill is itself negatively correlated with load in these two structures. Thus, NPFF marks neurons that track filling of the proximal duodenum.

To confirm that duodenal stimulation is sufficient to drive NPFF neuron activation, we next infused nutrient solutions directly into the duodenum via an intraduodenal (ID) catheter (Fig. 2K-M). We found that NPFF neurons were activated by ID infusion of nutrients (Intralipid: 1.30 ± 0.57 z, p = 0.029, glucose: 1.30 ± 0.23 z, p = 0.004, protein: 0.27 ± 0.40 z, p = 0.990 and mannitol: 1.34 ± 0.44 z, p = 0.026) but not saline (−0.27 ± 0.19 z), and the magnitude of these responses was similar to that of IG infusion of the same substance (IG, glucose: 1.64 ± 0.52 z, p = 0.026, intralipid: 0.64 ± 0.26 z, p = 0.030, protein: 0.77 ± 0.14 z, p = 0.0005 and mannitol: 2.26 ± 0.57 z, p = 0.012 vs saline: −0.27 ± 0.13 z). Notably, ID or IG mannitol elicited responses comparable to caloric nutrients, suggesting that NPFF neuron activation does not require calories and may instead reflect intestinal distension (which is induced by non-caloric mannitol and nutrient solutions but not saline).

Whereas IG infusion produced discrete, transient peaks in NPFF activity, ID infusion of the same solution produced sustained activation (Fig. 2D and 2L). This is consistent with the hypothesis that the pulsatile pattern during IG infusion reflects periodic boluses of nutrient emptying from the stomach. In support of this, we found that mimicking pulsatile emptying by delivering 0.1 mL boluses by ID infusion resulted in transient peaks in NPFF activity that mirrored the periodic activation following IG infusion (Fig. 2N-P). Moreover, NPFF neurons responded significantly faster to ID than IG infusion (median response onset: 22.1 s vs. 47.8 s, p = 0.0274, Fig. 2Q), which is consistent with a response to intestinal stimulation. Strikingly, the latency of the neural response to ID infusions (median response onset, 22.1 s, Fig. 2Q) was almost identical to that of the neural response to natural duodenal filling from the stomach, as measured by fluoroscopy (22.8 s, Fig. 2J). Taken together, these data show that NPFF neurons track the dynamics of filling and emptying of the proximal intestine.

One unusual aspect of cNTS dynamics is that many neurons respond differently to oral ingestion and IG infusion of the same food^41,42^. Since all the experiments above were performed by IG infusion, we wondered whether NPFF neurons also track duodenal fill during natural, oral ingestion. To test this, we gave animals intermittent access to a lick spout containing saline or Intralipid (Fig. S6), in a manner that matched our intermittent infusion experiments (Fig. 2N), and recorded NPFF neuron responses. We found that the onset of NPFF neuron activation after oral ingestion occurred with a latency of 26 ± 8 s from the first lick (Fig. S6D), which is consistent with both the general timescale of GI feedback^51^ and our specific results from infusion experiments (Fig. 2Q) but slower than typical orosensory responses^41,42^. Moreover, NPFF neuron activation continued to ramp well after the end of the one-minute period of access to the lick spout (peak response time = 180 ± 19 s). This is consistent with responses to duodenal filling as the stomach empties (which continues after ingestion stops) but not oral sensation. Thus, these results confirm that NPFF neurons track post-ingestive signals during natural ingestion.

### NPFF neurons trigger reciprocal changes in stomach and intestinal motility

We next investigated the function of these NPFF dynamics and asked whether activation of NPFF neurons alters GI function. To examine gastric emptying, we prepared mice for simultaneous optogenetic stimulation of NPFF neurons and X-ray fluoroscopy (Fig. 3A). Mice received an IG infusion (30% barium in water, 0.5 mL) along with simultaneous laser stimulation (20 Hz, 2s ON: 3s OFF), and stomach area was quantified over time. We found that in control animals (lacking ChR2 expression) stomach area gradually decreased over time as contents emptied into the intestine (Fig. 3A, bottom). In contrast, in animals expressing ChR2 that received stimulation, stomach area remained significantly elevated throughout the experiment (laser OFF: 28.2 ± 1.7 % vs laser ON: 58.9 ± 7.3 % of max at t = 40 min, p < 0.0001, Fig. 3B– D). This inhibition of gastric emptying was not due to changes in the rate of antral contractions (control: OFF: 9.27 ± 0.68 s vs. ON: 9.58 ± 0.50 s, p = 0.626; ChR2: OFF: 8.37 ± 0.74 s vs. ON: 8.35 ± 0.49 s, p = 0.376, Fig. 3E, Video S4). Moreover, we confirmed this result by showing that optogenetic stimulation reduced gastric emptying using an independent assay (acetaminophen test, Fig. 3F, G). Thus, these data support a model in which NPFF activation by duodenal filling feeds back to delay further emptying from the stomach.

**Figure 3.**
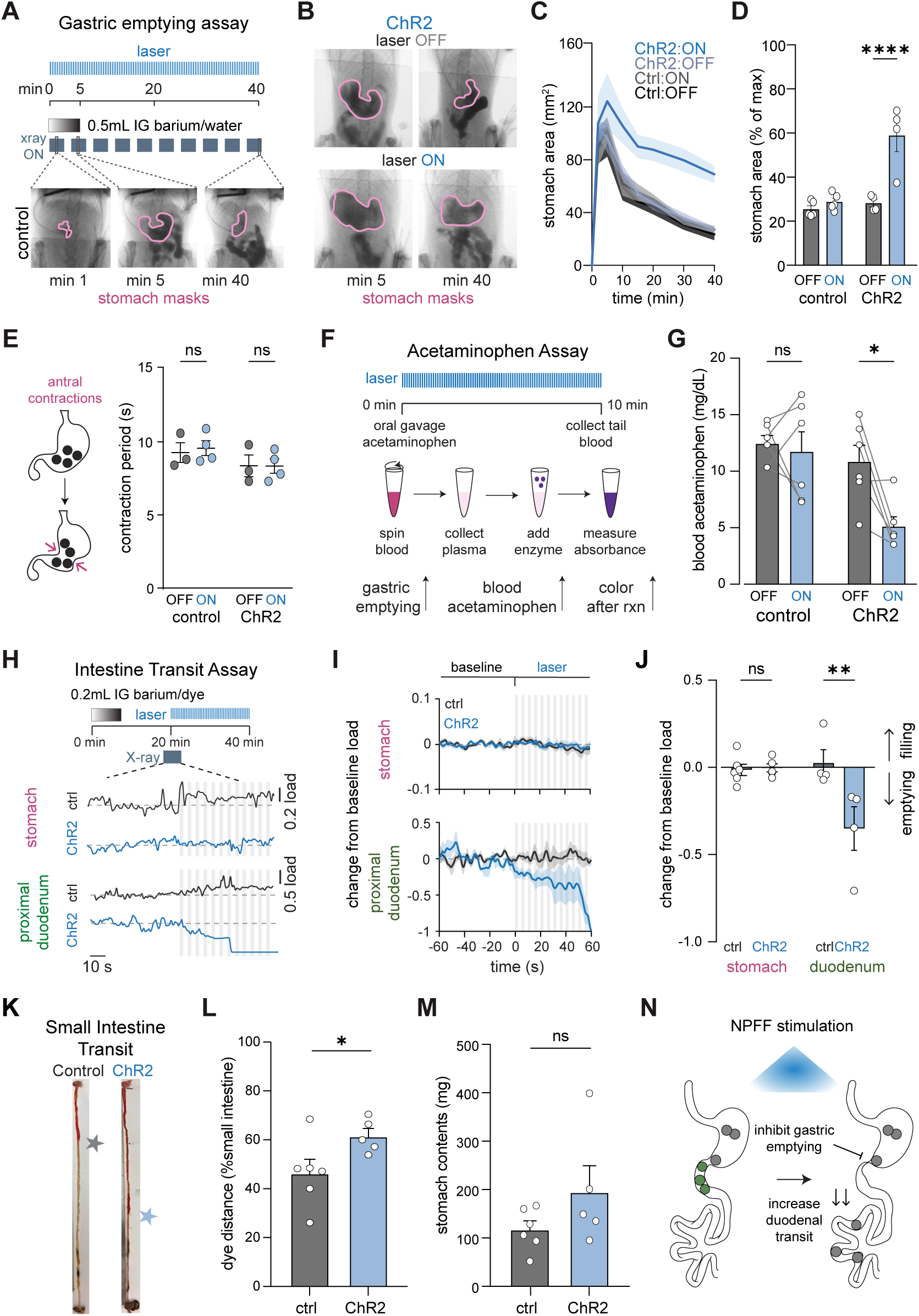
Stimulation of NPFF neurons inhibits gastric emptying while accelerating intestinal transit. **A.**Top, experimental timeline. *Npff^Cre^:: Ai32* (ChR2, n = 4) *or* littermate control (n = 5) mice were infused intragastrically with 0.5mL barium/water solution and imaged using an X-ray camera. Blue laser (20Hz, 2s ON: 3s OFF) was delivered for 40 min during ON condition. Bottom, representative fluoroscopy images from a control mouse t = 1, 5, and 40 min after infusion. **B.** Representative fluoroscopy images from a ChR2 mouse at t = 5 and 40 min during laser OFF (top) and ON (bottom) conditions. **C.** Stomach area over time in control and ChR2 mice during laser OFF vs ON. **D.** Stomach area at t = 40 min (% of max area) for control and ChR2 mice during laser OFF vs ON. **E.** Left, schematic illustrating antral contractions. Right, antral contraction period during laser OFF and ON conditions in control and ChR2 mice. **F.** Blood acetaminophen concentration was measured as a proxy for gastric emptying rate. **G.** Blood acetaminophen concentration at t = 10 min in control (n = 6) and ChR2 (n = 6) mice during laser OFF and ON conditions. **H.** Top, experimental timeline. ChR2 (n = 4) or control (n = 4-6) were infused intragastrically with 0.2 mL water/ barium solution containing red food dye. After 20 min, laser stimulation started and a fluoroscopy video was acquired 1 min before and after laser onset. After 20 min of stimulation, mice were perfused and intestines were dissected. Bottom, representative normalized load traces from a control (black) and ChR2 (blue) mouse showing stomach (top) and proximal duodenum (bottom) before and after laser onset. Load is normalized to the mean of the 60s pre-laser baseline period. Gray vertical lines indicate laser pulses. **I.** Average normalized stomach (top) and duodenum (bottom) load aligned to laser onset for control (black) and ChR2 (blue) mice. **J.** Change in normalized stomach and duodenum load from pre-laser baseline in control and ChR2 mice. **K.** Example intestines from a control and ChR2 mouse. Stars indicate the dye front. **L.** Distance traveled by red dye as a percentage of total small intestine distance in control and ChR2 mice. **M.** Stomach contents at the end of the intestine transit assay in control and ChR2 mice. Stimulation began only after gastric emptying was mostly complete (see Methods), so stomach contents did not differ between groups at this time point, confirming that the enhanced intestinal transit observed with ChR2 stimulation is not secondary to a difference in gastric emptying. **N.** Schematic illustrating the effects of NPFF stimulation on the gastrointestinal tract. Stimulation inhibits gastric emptying and increases duodenal transit. *p < 0.05, **p < 0.01, ****p < 0.0001, ns, not significant. Data are mean ± s.e.m. Each dot represents a single mouse.

Duodenal fill could be regulated entirely through changes in gastric emptying or it could involve coordinated changes in stomach and intestinal motility. To test this, mice received IG infusion of a small volume of barium (30% barium in water, 0.2 mL) and then were subjected to laser stimulation 20 minutes after infusion start, when most gastric contents had already emptied (Fig. 3H). We found by fluoroscopy that laser stimulation produced a rapid decrease in proximal duodenal load in ChR2 mice (ChR2: −0.351 ± 0.125 vs. ctrl: 0.026 ± 0.076 au, p = 0.0037, Video S5), whereas stomach load was unchanged (ChR2: −0.003 ± 0.023 vs ctrl: −0.014 ± 0.032 au, p = 0.99; Fig. 3I, J). This suggests that NPFF neuron stimulation accelerates movement of material out of the proximal duodenum.

To obtain an independent measure of the effect of NPFF stimulation on intestinal transit, we stimulated NPFF neurons for an additional 20 min in the experiment above and then extracted the intestines and quantified the total distance that the barium bolus traveled, which was visualized by a red dye. The dye traveled significantly further along the small intestine in ChR2 mice than in controls (ChR2: 61.6 ± 3.0 % vs ctrl: 46.4 ± 5.6 % of small intestine length, p = 0.046; Fig. 3K-N), confirming that NPFF neuron activation accelerates intestinal transit.

To directly confirm that NPFF neuron stimulation drives proximal intestinal motor activity, we developed a preparation in which we could record electromyogram (EMG) activity throughout the GI tract while optogenetically stimulating NPFF neurons in anesthetized mice (Fig. 4A, B). We found that laser stimulation of ChR2-expressing mice caused a sharp increase in the spike rate in the proximal duodenum (pre: 0.3 ± 0.1 vs. during: 6.8 ± 2.1 spikes/(min*electrode), p = 0.013) within 25 ± 7 s and this remained elevated above baseline following laser offset (post: 2.0 ± 0.5 spikes/(min*electrode), p = 0.01, Fig. 4C,D, Video S6). This confirms that NPFF neuron activity controls intestinal motor output.

**Figure 4.**
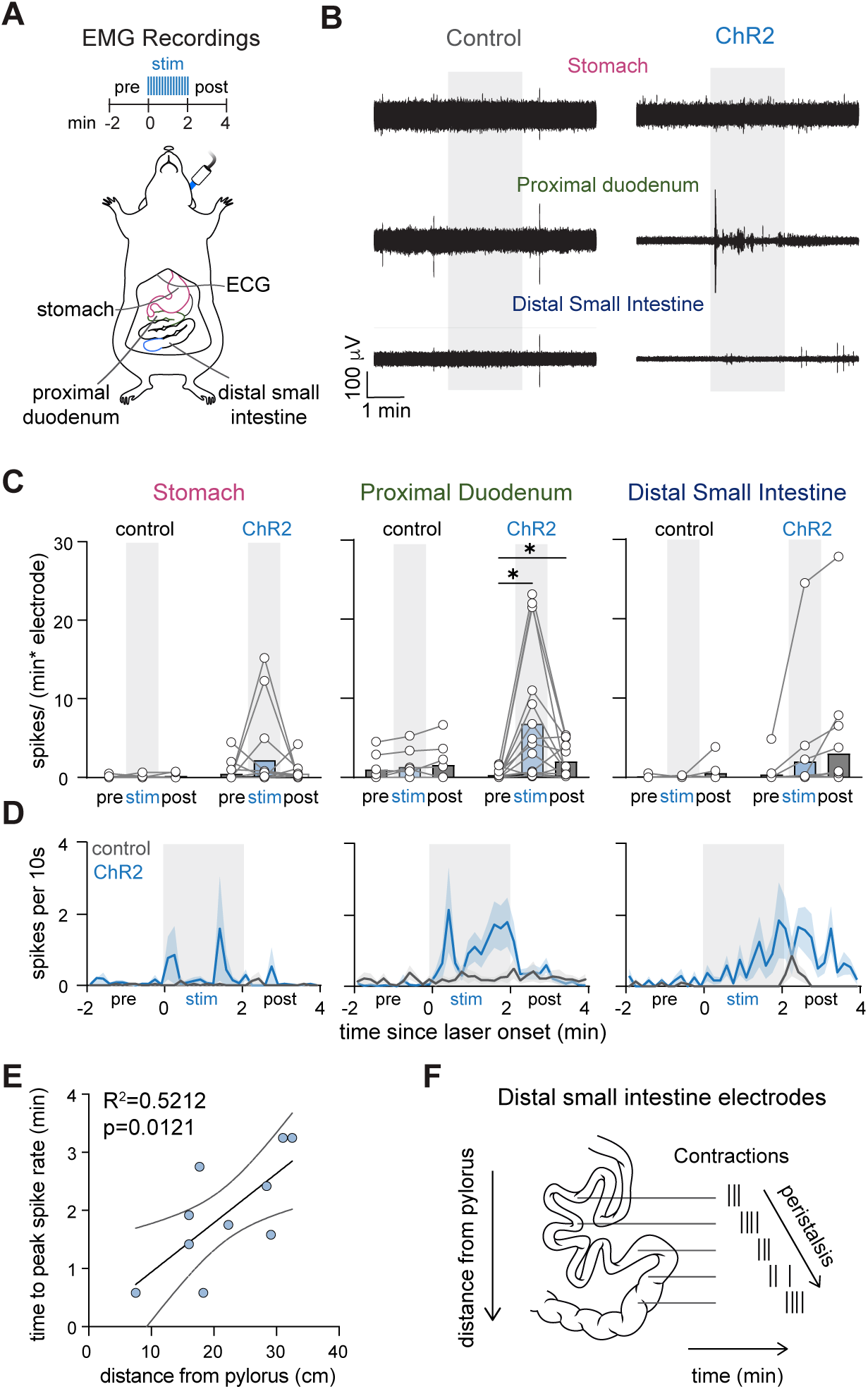
Stimulation of NPFF neurons drives motor activity in the proximal duodenum. **A.** Schematic of EMG recording setup during NPFF optogenetic stimulation. Electrodes were implanted on the stomach, proximal duodenum, and distal small intestine, with simultaneous ECG recording. *Npff^Cre^:: Ai32* (ChR2) or control littermate mice received blue laser stimulation via an optical fiber targeting the NTS. **B.** Representative EMG traces from stomach, proximal duodenum, and distal small intestine electrodes in a ChR2 (top) and control (bottom) mouse during laser stimulation (gray). Scale bar, 100 µV, 1 min. **C.** EMG spike rate (spikes/ (min*electrode)) in the stomach, proximal duodenum, and distal small intestine pre, during and post laser stimulation in control (n = 8, gray) and ChR2 (n = 16, blue) mice. **D.** Mean EMG spike rate (spikes/10s) aligned to laser onset for stomach, proximal duodenum, and distal small intestine in control (gray) and ChR2 (blue) mice. Gray shading indicates laser ON period. **E.** Linear regression of time to maximal EMG spike rate following laser onset vs. distance of distal electrodes from the pylorus. Each dot represents a single electrode. **F.** Schematic showing contractions traveling distally across electrodes during peristalsis. *p < 0.05. Each dot represents a single mouse unless specified.

In contrast to the proximal duodenum, there was no significant change in stomach or distal small intestine spike rate during stimulation (all ns; Fig. 4C, D). However, we noticed visually that contractions propagated distally along the intestine and observed a clear trend in which the spike rate in the distal intestine appeared to ramp as stimulation progressed (pre: 0.3 ± 0.3 vs during: 2.0 ± 1.5 vs post: 3.0 ± 1.8 spikes/(min*electrode), Fig. 4C, D). To test whether activity initiated in the proximal duodenum propagates distally with a delay, we correlated the latency to the peak in spiking following laser stimulation with the distance from the pylorus of individual electrodes in the distal intestine. This revealed that the time to maximal spike rate following laser onset increased linearly with electrode distance from the pylorus (R² = 0.52, p = 0.012; Fig. 4E-F), consistent with peristaltic activity initiating in the proximal duodenum and propagating distally through local mechanisms^17,19,20^. These data demonstrate that NPFF neuron activation induces opposing responses in the stomach and intestine that function, together, to pace the flow of nutrients through the GI tract.

### NPFF neurons inhibit feeding through a cholinergic mechanism

Because NPFF neuron activation altered GI motility, we next asked whether activation of this circuit influences feeding behavior. Mice expressing ChR2 in NPFF neurons were fasted overnight and then allowed to refeed for one hour (Fig. 5A). We found that optogenetic stimulation reduced intake of both chow (laser OFF: 0.52 ± 0.02 g vs. laser ON: 0.41 ± 0.04 g, p = 0.048) and high-fat diet (HFD, laser OFF: 0.96 ± 0.12 g vs. laser ON: 0.36 ± 0.11 g, p = 0.0035) in ChR2 mice, whereas control animals that lack ChR2 expression were unaffected (Fig. 5B). This indicates that NPFF neuron activation inhibits solid food consumption.

**Figure 5.**
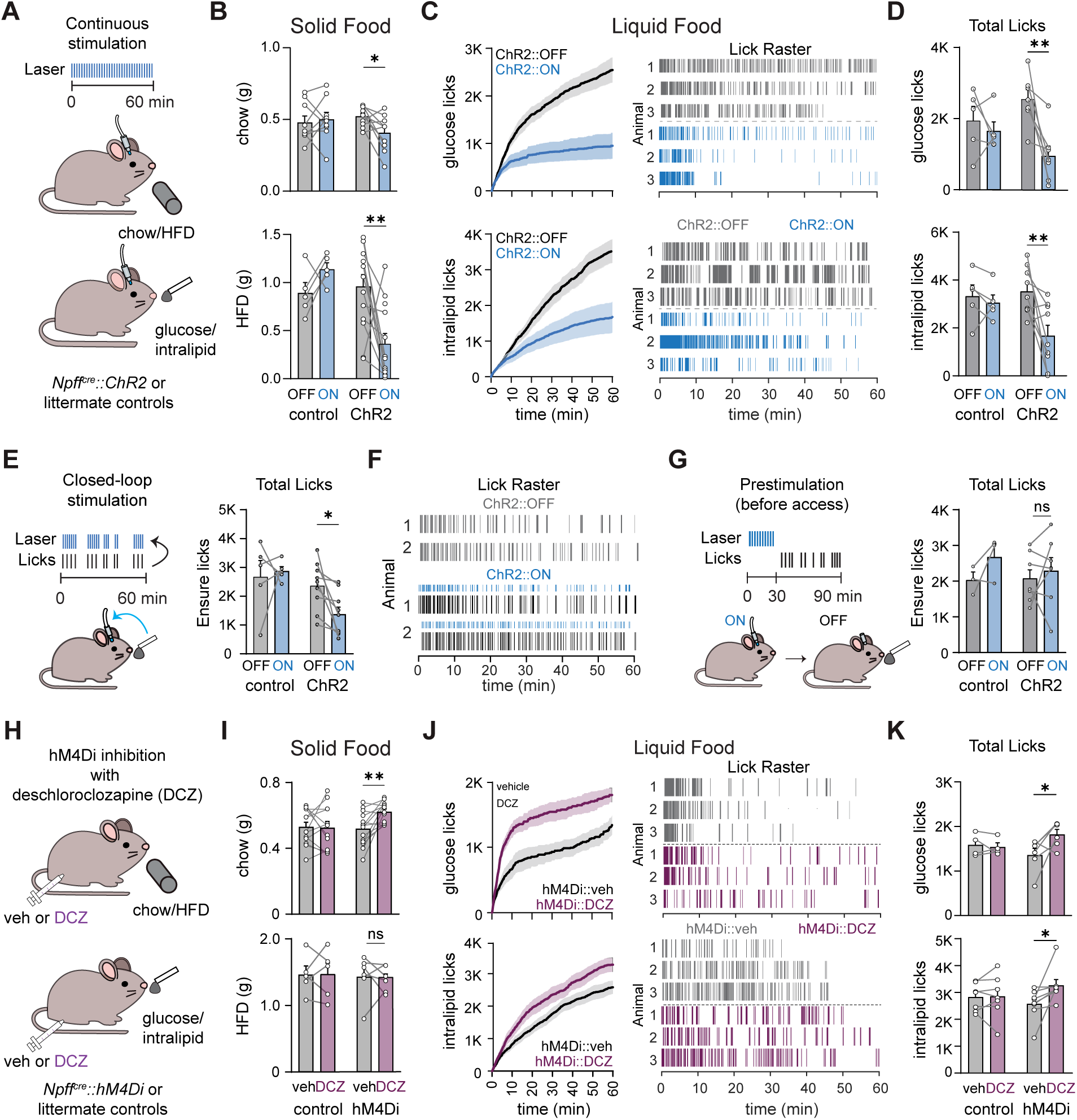
NPFF neurons bidirectionally regulate food intake. **A.** Optogenetic stimulation of NPFF neurons. *Npff^Cre^::Ai32* (ChR2) and littermate control mice were implanted with an optical fiber targeting the NTS and given access to chow, high fat diet (HFD; top, solid), glucose, or intralipid (bottom, liquid). Blue laser (20Hz, 2s ON: 3s OFF) was delivered for 60 min during ON condition. **B.** Chow (top) and HFD (bottom) consumption (g) during a 1-h fast-refeed in control and ChR2 mice (chow: n = 9 ctrl, n = 10 ChR2; HFD: n = 5 ctrl, n = 13 ChR2). **C.** Left, cumulative licks over time during glucose (top) and intralipid (bottom) consumption in control and ChR2 mice (glucose: n = 5 ctrl, n = 7 ChR2; intralipid: n = 5 ctrl, n = 9 ChR2). Right, example lick rasters from 3 ChR2 mice during laser OFF (gray) and ON (blue) conditions. Each tick indicates a lick. **D.** Total licks during glucose (top) and intralipid (bottom) consumption. **E.** Left, schematic for closed loop stimulation: laser is triggered by licking over a 60 min session. Right, total Ensure licks with laser off vs. on in control and ChR2 mice (n = 5 ctrl, n = 9 ChR2). **F.** Example lick rasters from 2 ChR2 mice during laser OFF (gray) and ON (black) conditions. Blue ticks indicate when the laser was triggered. **G.** Left, schematic for prestimulation: NPFF neurons are stimulated for 30 min then mice are given food access for 60 min. Right, total Ensure licks in control and ChR2 mice (n = 3 ctrl, n = 7 ChR2). **H.** Chemogenetic inhibition of NPFF neurons. *Npff^Cre^::hM4Di* (hM4Di) or littermate control mice were injected with vehicle or deschloroclozapine (DCZ) and given access to chow, HFD (top), glucose, or intralipid (bottom). **I.** Chow (top) and HFD (bottom) consumption (g) during a 1h fast-refeed in control and hM4Di mice (chow: n = 13 ctrl, n = 14 hM4Di; HFD: n = 5 ctrl, n = 6 hM4Di). **J.** Left, cumulative licks over time during glucose (top) and intralipid (bottom) consumption in control and hM4Di mice (glucose: n = 4 ctrl, n = 6 hM4Di; intralipid: n = 7 ctrl, n = 8 hM4Di). Right, example lick rasters from 3 hM4Di mice during vehicle (gray) and DCZ (purple) conditions. Each tick indicates a lick. **K.** Total licks during glucose (top) and intralipid (bottom) consumption. *p < 0.05, **p < 0.01, ns, not significant. Data are mean ± s.e.m. Each dot represents a single mouse.

To better understand how NPFF neurons influence the dynamics of food intake, we also measured consumption of glucose and Intralipid solutions. Optogenetic stimulation reduced total licks in ChR2 mice for both glucose (OFF: 2544 ± 267 vs. ON: 949 ± 280, p = 0.0056) and Intralipid (OFF: 3522 ± 327 vs. ON: 1675 ± 436, p = 0.0064), with no effect in controls (Fig. 5C-D). Analysis of feeding microstructure revealed that stimulation had nutrient-specific effects on bout structure (Fig. S8). These effects on behavior were time locked to laser stimulation, since prestimulation of NPFF neurons for 30 min before food access had no effect on total intake, whereas closed-loop (lick-triggered) stimulation effectively reduced consumption (Fig. 5E-G).

To test whether NPFF activity is necessary for regulation of food intake, we next silenced these neurons and measured consumption of different diets (Fig. 5H). We found that chemogenetic silencing of NPFF neurons with hM4Di increased consumption of chow (ctrl veh: 0.53 ± 0.03 vs. deschloroclozapine, DCZ: 0.52 ± 0.04, p = 0.987 and hM4Di veh: 0.52 ± 0.03 vs. DCZ: 0.62 ± 0.02 g, p = 0.005) but not HFD, which may reflect a ceiling effect, as animals consumed much more HFD (Fig. 5I). Silencing also increased consumption of liquid diets glucose (ctrl veh: 1584 ± 128 vs. DCZ: 1535 ± 97, p = 0.959 and hM4Di veh: 1356 ± 149 vs. DCZ: 1819 ± 114 licks, p = 0.030) and Intralipid (ctrl veh: 2840 ± 197 vs. DCZ: 2865 ± 309, p = 0.99 and hM4Di veh: 2582 ± 204 vs. DCZ: 3277 ± 221 licks, p = 0.018, Fig. 5J, K), and these effects were mediated by an increase in bout size, rather than bout number (Fig. S8). Together, these data indicate that the natural activation of NPFF neurons by duodenal fill plays a physiologic role in controlling ingestive behavior.

Given that NPFF neurons are activated by GI signals (Fig. 2) and feed back to alter GI motility (Fig. 3, 4), it is possible that NPFF neurons inhibit food intake via their effects on the gut. However, it is also possible that NPFF neuron activity has a direct effect on behavior. To investigate possible circuit mechanisms, we expressed ChR2-eYFP in these cells and visualized axonal projections throughout the brain (Fig. 6A). We observed dense terminal fields in the dorsal motor nucleus of the vagus (DMV) and the parabrachial nucleus (PBN), but not other brain regions, suggesting these are the primary output targets^52^ (Fig. 6B). To determine the functional significance of these and other projections, we stimulated NPFF neurons expressing ChR2 and then quantified FOS expression in these regions (Fig. 6C). This revealed dramatic FOS induction in the DMV (ctrl: 165 ± 14 vs. ChR2: 394 ± 36 cells/mm², p = 0.0001) as well as the subpostrema itself (ctrl: 68 ± 27 vs. ChR2: 328 ± 64 cells/mm², p < 0.0001) but surprisingly no significant increase in FOS expression in the PBN (Fig. 6D). Importantly, we confirmed that many of the activated neurons in the DMV are peripherally projecting motor neurons (in ChR2 animals, 56 ± 11% Fos+ cells were also Fluorogold+ and 22 ± 3% of all Fluorogold+ cells were also Fos+, Fig. 6E). Together, these findings identify the DMV, but not the PBN, as a primary functional output target of NPFF neurons.

**Figure 6.**
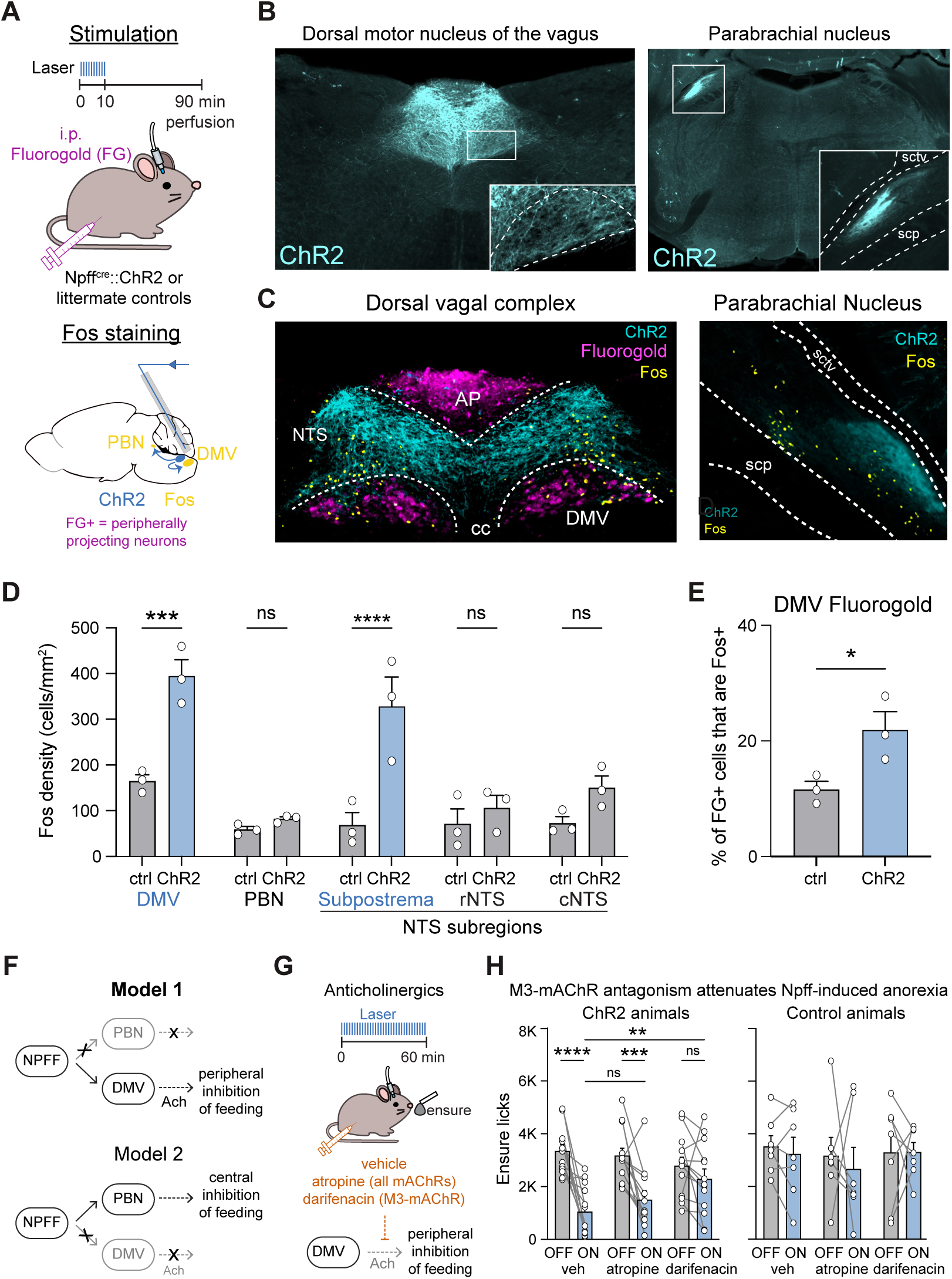
NPFF neurons inhibit feeding through a cholinergic mechanism. **A.** ChR2-assisted Fos mapping. *Npff^Cre^::Ai32* (n=3) and control (n=3) mice were injected intraperitoneally with Fluorogold (FG) to label peripherally projecting neurons, stimulated for 10 min, and perfused 90 min after stimulation onset for Fos immunostaining. **B.** Representative images of ChR2-expressing NPFF neuron terminals in the dorsal motor nucleus of the vagus (DMV, left) and parabrachial nucleus (PBN, right). Insets show zoomed views of terminal fields. sctv, ventral spinocerebellar tract; scp, superior cerebellar peduncle. **C.** Representative coronal sections showing ChR2 terminals (cyan) and Fos expression (yellow) in the dorsal vagal complex (left) and parabrachial nucleus (right). Fluorogold (magenta) labels the AP and DMV. NTS, nucleus of the solitary tract; cc, central canal. **D.** Fos density (cells/mm²) in the DMV, PBN, subpostrema, rNTS, and cNTS in control and ChR2 mice. Subpostrema is a custom border region combining the area postrema and NTS atlas borders. **E.** Percentage of FG+ cells that are also Fos+ in the DMV in control (gray) and ChR2 (blue) mice. **F.** Schematic of two candidate models for how NPFF neuron stimulation suppresses feeding: peripheral inhibition via a cholinergic DMV projection (Model 1) versus central inhibition via a PBN projection (Model 2). **G.** Schematic of pharmacological experiment testing whether cholinergic signaling downstream of the DMV mediates NPFF-induced feeding suppression. *Npff^Cre^::Ai32* (ChR2, n= 13) and control (n = 7) mice received vehicle, methylatropine (broad muscarinic acetylcholine receptor antagonist), or darifenacin (M3 muscarinic receptor antagonist) prior to laser stimulation and were given access to Ensure for an hour. **H.** Left, total ensure licks after vehicle, atropine or darifenacin injection in ChR2 mice. Right, same in controls. *p < 0.05, **p < 0.01, ***p < 0.001, ****p < 0.0001, ns, not significant. Data are mean ± s.e.m. Each dot represents a single mouse.

DMV neurons are cholinergic, and their effects on gut motility are mediated by the peripheral release of acetylcholine^29,30^ (Fig. 6F). We therefore reasoned that if NPFF neurons inhibit food intake through these descending vagal motor circuits, then blockade of acetylcholine signaling in the periphery should prevent anorexia induced by NPFF activation (Fig. 6G). To this end, we used two drugs that prevent signaling through muscarinic acetylcholine receptors and have poor brain penetration: methylatropine, which blocks all muscarinic receptors, and darifenacin^53,54^, which selectively targets the M3 isoform that is expressed in smooth muscle and is critical for GI motility^55–59^. In the absence of drug, we found that optogenetic stimulation of NPFF neurons strongly decreased Ensure intake (Fig. 6H). Strikingly, pretreatment with darifenacin abolished this optogenetically-induced suppression of feeding, indicating that peripheral M3 receptor activity is required for NPFF food intake suppression (Fig. 6H). Atropine, on the other hand, did not block the anorexia caused by NPFF stimulation, possibly because it also blocks other muscarinic receptors whose antagonism can enhance cholinergic transmission and offset the effect of M3 blockade^58,60,61^. Importantly, neither drug had any effect on food intake at baseline, indicating that these effects are not due to a non-specific change in total consumption (Fig. 6H).

This provides evidence that NPFF neurons inhibit food intake through their effects on the GI tract. More broadly, the data presented here show that NPFF neurons, in response to activation by duodenal fill, trigger three complementary responses to ensure the appropriate flow of nutrients through the gut: suppression of food intake, inhibition of gastric emptying and acceleration of intestinal transit.

## Discussion

Gastrointestinal function is controlled by combination of top-down signals from the brain and local circuits in the gut^19,30,62^. A role for the brain was established as early as 1898, when Cannon showed that stress can suppress gastric contractions^45,63^. A year later, Bayliss and Starling found that peristaltic contractions persist in the absence of extrinsic innervation, indicating a role for autonomous local circuits^20^. While a tremendous amount has been learned in the intervening century, it still remains debated how exactly these intrinsic and extrinsic neural circuits work together to enable the choreographed movement of food through the GI tract^24,64^.

We reasoned that we could obtain new insight into this old question by combining X-ray fluoroscopy, which provides real-time information about the flow of food through the gut, and calcium recordings in the caudal brainstem, which provide real-time information about the dynamics of the neurons that receive direct input from the GI tract. Using this approach, we showed that the proximal duodenum fills and empties in discrete pulses, and that a specific population of cNTS neurons defined by expression of NPFF tracks these pulsatile dynamics. We showed that stimulation of NPFF neurons initiates a coordinated motor program that inhibits gastric emptying while accelerating intestinal transit, thereby clearing the proximal duodenum for the next bolus of food. Moreover, we showed NPFF neurons bidirectionally control food intake via these GI motor effects. This reveals a cell type in the brainstem that orchestrates the pacing of food transit (Fig. 7).

**Figure 7.**
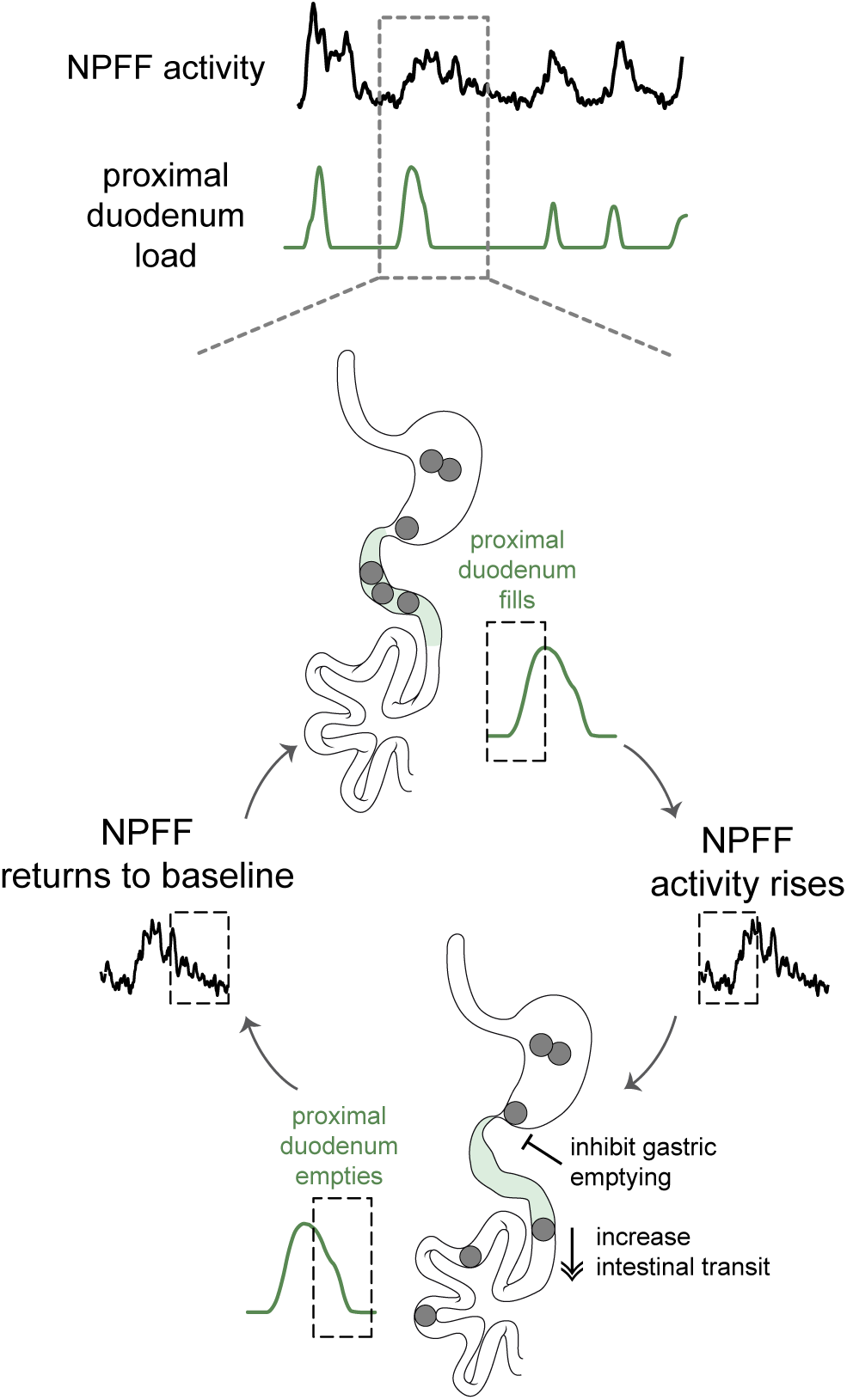
NPFF neurons form a closed feedback loop that paces the flow of food through the proximal intestine. Top, schematized recordings of NPFF neuron activity (fiber photometry, black) and proximal duodenal content (fluoroscopy-derived load, green) illustrate the periodic, coupled dynamics of the two signals over the course of a session. Bottom, the filling of proximal duodenum (shaded in green) activates NPFF neurons in the subpostrema, which inhibit gastric emptying and accelerate intestinal transit, together clearing the duodenum. As duodenal content falls, NPFF activity returns to baseline, resetting the circuit until the next filling event initiates the next cycle.

### The neural substrates of intestinal brakes are largely unknown

It is well established that the entry of food into different segments of the intestine can trigger inhibition of gastric emptying and suppression of food intake. These reflexes, which are known as “brakes” (e.g. duodenal brake^10,26,65–67^ and ileal brake^68–71^), function to prevent intestinal overfilling and have been characterized using a variety of pharmacological and surgical approaches. While some of the relevant hormones and afferent nerves have been identified, we know essentially nothing about the cell types in the brain that mediate these classic reflexes. Of note, although several cell types can inhibit gastric emptying when artificially stimulated, none of these cells have been shown to track any aspect of natural food transit, and many respond instead to other signals such as stress, inflammation, emetics, or hormones^51,72–80^. Moreover, while the cNTS is an essential node in the reflex loops controlling intestinal brakes^24,49^, recent single-cell imaging revealed that most cNTS cells respond, counterintuitively, to rapid orosensory feedback rather than slower GI cues^41,42^. This has raised the question of which cells in the caudal brainstem are responsible for these fundamental reflexes.

We reasoned that cNTS neurons responding to duodenal stimuli are likely to be located medially in the vicinity of the subpostrema, based on both responses from anesthetized recordings^31,81^ and the anatomy of the central terminals of vagal cell types innervating the intestine^32^. Expression of NPFF identifies a unique cNTS cell type that is highly restricted to the subpostrema^34,50,52,82,83^ (Fig. S3) and, consistently, we showed that it responds to duodenal filling. Stimulation of NPFF neurons inhibited gastric emptying, as expected for a cell type that participates in an intestinal brake, but surprisingly also accelerated intestinal transit. This is noteworthy because brakes have traditionally been described solely as inhibitory reflexes in which nutrients in the intestine act to arrest further nutrient delivery from upstream. Our data indicate that this is only half of the mechanism, and that the same neurons that stop gastric emptying simultaneously accelerate intestinal motility. These two actions are complementary and coupling them to a single cell type ensures that the two organs are always driven in the appropriate relationship to one another. This suggests that the GI reflexes that have been described as brakes are better thought of as push-pull mechanisms for pacing.

### NPFF neurons are an entry point into the mechanisms controlling GI motility

Multiple aspects of GI motility can be modulated by the central nervous system^84^. For the stomach, this includes accommodation (which involves an increase in size of the fundus) as well as mixing and emptying (which are controlled by the antrum and pyloric sphincter)^5,25,29,45,85,86^. Stimulation of NPFF neurons inhibits gastric emptying without altering the rate of antral contractions, indicating that this effect is likely mediated by control of the pyloric sphincter^87–89^. Of note, closing the pylorus while preserving antral contractions promotes mixing of gastric contents^90^, indicating this is one function of NPFF neurons. On the other hand, we observed no gross effect of NPFF neuron stimulation on fundal size, which is consistent with the idea that these neurons are not involved in promoting accommodation. This is not the case for all cell types in this brain region, as we have recently shown that stimulation of adjacent neurons in the area postrema (GFRAL neurons) causes dramatic fundal expansion^51^. This demonstrates that different molecularly defined cell types in the caudal brainstem are coupled to different GI reflexes.

In contrast to the stomach, much less is known about how the caudal brainstem modulates intestinal motility. A few studies have shown that hormone injection into the brain can increase intestinal transit^91–93^, but little is known about specific cells or circuits. We found that NPFF neurons specifically increase motility in the proximal duodenum, which then propagates to the distal intestine, presumably through intrinsic circuits. Consistent with this, the DMV innervation of the intestine is most dense in the proximal duodenum and then greatly declines distally^88,94^.

There is a paucity of information about the local circuitry in the caudal brainstem that connects the NTS to the DMV to control gut motility. Most of our knowledge is based on experiments in which drugs blocking different neurotransmitter systems were microinjected into the DMV of anesthetized animals^29,76^. This has led to a model in which the key projection from the NTS to the DMV is GABAergic and provides tonic inhibition that suppresses gastric contractions^23,29^. However, local pharmacology cannot provide information about specific cell types or interconnections, and studies in anesthetized animals do not measure the natural flow of food through the gut^95^. In contrast to this earlier work, our studies identified a glutamatergic cell type (NPFF neurons) that senses and responds to food to control motility of both the stomach and intestine. This revises current models for how cNTS circuits control GI motility, although it is important to note that additional cell types are certainly involved. Going forward, there is an opportunity to use the strategy we have described here – which is based on the simultaneous imaging of the gut and brainstem in awake animals – to systematically map these gut-brain circuits that control digestion.

## Methods

All experimental protocols were approved by the Institutional Animal Care and Use Committee of the University of California, San Francisco, following the National Institutes of Health guidelines for the Care and Use of Laboratory Animals.

### Mouse strains

Experimental animals (>8 weeks old, both sexes) were kept in temperature- and humidity-controlled facilities on a 12-h light-dark cycle with ad libitum access to water and standard chow (PicoLab 5053). The following mice were obtained from the Jackson Laboratory: WT (C57BL/6J; 000664), Gpr65-ires-Cre (Gpr65^tm1.1(cre)Lbrl^/RcngJ, 029282), Ai162 (B6.Cg-Igs7^tm162.1(tetO-GCaMP6s,CAG-tTA2)Hze^/J, 022731), TIGRE2-jGCaMP8s-IRES-tTA2 (Igs7tm2^(tetO-GCaMP8s,CAG-tTA2) Genie/^J, 037719), Ai32 (B6.Cg-Gt(ROSA)^26Sortm32(CAG-COP4*H134R/EYFP)Hze^/J, 024109), R26-LSL-Gi-DREADD (B6.129-Gt (ROSA)26Sor^tm1(CAG-CHRM4*,-mCitrine)Ute^/J, 026219) and Ai14 (B6.Cg-Gt(ROSA)26Sor^tm14(CAG-tdTomato)Hze^/J; 007914).

Npff-2A-Cre mice were generated by homologous recombination at the endogenous *Npff* locus, aided by targeted CRISPR endonuclease activity. The targeting vector was constructed containing a T2A-Cre cassette inserted immediately upstream of the endogenous stop codon, a 2 kb upstream homology arm, and a 1kb downstream homology arm. An sgRNA was selected (CTTCCCAAAGCGCTGAGGAG) to introduce CRISPR double strand breaks near the stop codon, and the corresponding PAM sequence was mutated in the targeting vector (CGG to CTG) to prevent vector cleavage. Super-ovulated female FVB/N mice were mated to FVB/N stud males, and fertilized zygotes were collected from oviducts. Cas9 protein (100 ng/uL), sgRNA (250 ng/uL), and targeting vector DNA (100 ng/mL) were mixed and injected into the pronucleus of fertilized zygotes. Zygotes were implanted into oviducts of pseudopregnant CD1 female mice. Screening of the pups by qPCR identified independent founder lines that contained insertion of Cre but not sequences from the targeting vector (i.e. knock-ins). These founders were crossed to Ai14 reporter mice, and a line that showed a brain-wide recombination pattern that was identical to previous reports of *Npff* expression (i.e. restricted to the cNTS in the vicinity of the subpostrema) was selected to expand and further characterized by *in* situ hybridization for endogenous *Npff* expression (Fig. S4).

Npff-2A-Cre mice were crossed to jGCaMP8s-IRES-tTA2, Ai32, R26-LSL-Gi-DREADD, and Ai14 mice to generate *Npff-Cre/+::GCaMP8s/+, Npff-Cre/+::ChR2/+, Npff-Cre/+::hM4Di/+,* and *Npff-Cre/+::tdTomato/+ mice,* respectively. Afferents from Gpr65 neurons in the nodose that project to the NTS were visualized by crossing Gpr65-Cre/+ and Ai162.

### Intracranial surgery

#### General procedures

Animals were anesthetized with 2% isoflurane and secured in a stereotaxic frame on a heating pad. Ophthalmic ointment was applied to protect the eyes, and subcutaneous injections of meloxicam (5.0 mg/kg) and sustained-release buprenorphine (1.5 mg/kg) were administered prior to surgery. The scalp was shaved and scrubbed with betadine and alcohol (three times), local anesthetic was applied (bupivacaine 0.25%), and a midline incision was made. A small craniotomy was performed using a dental drill (0.5 mm burr hole). Fiber optic cannulas were then implanted and secured to the skull with Metabond (Patterson Dental Supply, 07-5533559, 07-5533500; Henry Schein, 1864477) and Flow-It (Patterson Dental Supply, 07-6472542). To reduce postoperative inflammation, animals received dexamethasone (0.6 mg/kg) for three days following surgery.

#### Fiber photometry implants in the subpostrema

*Npff-Cre/+:: GCaMP8s/+* mice were prepared for photometry recordings by implanting an optic fiber (Doric Lenses, MFC_400/430-0.48_6mm_MF2.5_FLT) and sleeve (Doric Lenses, SLEEVE_BR_2.5) above the subpostrema (+1.4 mm anterior–posterior (AP), 0.1 mm medial–lateral (ML), and −4.1 mm dorsal–ventral (DV), relative to the occipital crest, at a 20° angle in the anterior-posterior direction). Mice were allowed to recover for at least 2 weeks before the first photometry experiment. For fluoroscopy experiments, head bars were affixed to the skull using Metabond and intragastric or intraduodenal catheters were placed in a separate surgery.

#### Optogenetic implants in the subpostrema

*Npff-Cre/+::ChR2/+* mice were prepared for optogenetic experiments by implanting a fiber optic cannula (Doric, MFC_200/240-0.22_6mm_ZF1.25(G)_FLT) above the subpostrema (+1.3 mm anterior–posterior (AP), 0 mm medial–lateral (ML), and −3.8 mm dorsal–ventral (DV) relative to the occipital crest, at a 20° angle in the anterior-posterior direction). Mice were allowed to recover for a minimum of 3 weeks before optogenetic experiments. For fluoroscopy experiments, head bars were affixed to the skull using Metabond and intragastric catheters were placed in a separate surgery.

### Intragastric and intraduodenal catheter surgery

#### General procedures

Animals were anesthetized with 2% isoflurane, and surgical sites were shaved and sterilized with alternating betadine and ethanol scrubs. Meloxicam (5 mg/kg) and sustained-release buprenorphine (1.5 mg/kg) were administered subcutaneously prior to surgery. A midline abdominal incision of approximately 1.5 cm was made caudal to the xiphoid process, and a secondary 1 cm incision was made between the scapulae for catheter externalization. Blunt dissection was used to separate the skin from the subcutaneous tissue along the left flank, creating a tunnel between the two incisions for catheter routing. A small incision was made in the abdominal wall, and the catheter (Instech, C30PU-RGA1439) was guided through the interscapular incision into the abdominal cavity using curved hemostats. The abdominal cavity was irrigated with 1 mL of sterile saline following placement (described separately below), and the abdominal wall was closed in two layers. The catheter was anchored to the muscle at the interscapular site, and the interscapular incision was closed. The external catheter end was capped with a 22-gauge PinPort (Instech, PNP3F22). Mice received Baytril (5 mg/kg) and warm saline at the end of surgery and were allowed to recover for two weeks before the start of experiments.

#### Intragastric catheter placement

The stomach was gently externalized with atraumatic forceps, and a purse-string suture was placed in the forestomach using 7-0 non-absorbable Ethilon. A small puncture was made at the center of the purse string, the catheter tip was inserted, and the suture was tightened to secure it in place, with 2–5 mm of catheter fixed within the stomach lumen. The stomach was then returned to its anatomical position.

#### Intraduodenal catheter placement

The proximal duodenum was gently externalized with atraumatic forceps and a purse-string suture was placed approximately 1–2 cm distal to the pyloric sphincter using 7-0 non-absorbable Ethilon. A small puncture was made at the center of the purse string, the catheter tip was inserted into the duodenal lumen, and the suture was tightened to secure it in place. The duodenum was then returned to its anatomical position.

### Fluoroscopy experiments

#### Fluoroscopy setup

Fluoroscopy recordings were done using a custom-built low energy fluoroscope (Glenbrook Technologies, Newark, NJ). Fluoroscopy data were acquired at 30 frames/s with a voltage of 25 kV and a current of 0.216 mA. Mice were habituated to head-fixation and body restraint in a custom conical tube for at least five days prior to the first experiment. During habituation, mice received IG infusions of water, Ensure, or Intralipid with and without barium. To minimize radiation exposure to mice, X-ray exposure was intermittent, lasting no more than 120 continuous seconds and no more than 5 minutes per session.

For the IG infusions, either water, 20% Intralipid (Sigma, I141-100ML; Medline, BHL2B6064H) or 24% glucose was mixed with the oral contrast barium suspension (barium sulfate oral suspension 60% w/v, Liquid E-Z-PAQUE, Bracco Diagnostics) for visualization under X-ray imaging, in a ratio of 1:1. Infusions were delivered using a syringe pump (Harvard Apparatus, 70-2001). The IG catheter was attached to the syringe pump using plastic tubing and adapters (AAD04119, Tygon; LS20, Instech).

#### Experimental approach

For all experiments, mice were head-fixed and fasted 3-4 hours prior to the start of the experiment to allow clearing of gastric contents.

##### GI dynamics characterization

Mice received an IG infusion of 0.5 mL barium/intralipid or barium/glucose over 5 min. Intralipid was chosen for the main analysis due to its macronutrient-dense composition and ease of visualization under X-ray. Ten 30-second video clips were acquired, distributed evenly across the recording session.

##### Simultaneous photometry and fluoroscopy

Mice received an IG infusion of 0.2 mL barium/intralipid or barium/glucose over 1 min. Fluoroscopy video was acquired intermittently in 30s-on 30s-off blocks over 10 min.

#### Analysis

##### Segmentation with SAM2

Raw fluoroscopy videos were separated into active clips, trimmed at both ends to exclude transition frames, and segments below a minimum length were discarded. Valid clips were then downsampled to 4 Hz and cropped to the region of interest. Cropped and downsampled clips were preprocessed with contrast-limited adaptive histogram equalization (CLAHE), non-local means denoising, and spatial/temporal despeckling to enhance visibility of the contrast agent.

Preprocessed videos were segmented using SAM2 (Segment Anything Model 2)^96^. Each organ, stomach, proximal duodenum, or distal small intestine, was segmented by a polygon mask drawn around the organ of interest on key frames spaced approximately every 50–100 frames, with the interval varying between animals depending on the amount of motion and image quality. The distal small intestine mask was placed on a segment clearly distal to the proximal duodenum and served as a control to confirm genuine GI motility and to eliminate global motion artifacts. SAM2 propagated these seeded masks across all frames of the clip. SAM2-derived masks were quality controlled using a custom MATLAB pipeline. Every frame was manually reviewed against the source video, and frames in which a mask was missing were corrected by applying the nearest temporally adjacent accepted mask. Finally, mask area and mean pixel intensity within the mask were computed for every frame.

##### Organ load quantification

Organ load was computed as area × (I_background_ − I_mask_), where I_background_ is a fixed background intensity value obtained by manually drawing a region of interest over an empty area of the fluoroscopy frame and averaging pixel intensity over the first several seconds of video, and I_mask_ is the mean pixel intensity within the SAM2-derived, QC-corrected mask.

For GI dynamics characterization and simultaneous photometry-fluoroscopy, load traces were normalized to each region’s own maximum value within the session (0–1 scale). Where a recording session comprised multiple discontinuous clips, a smoothed curve interpolated across the gaps between clips was used for visualization only; all statistical comparisons and correlations were computed using only the real, clip-derived data points.

##### Interpeak interval (IPI) analysis

From the normalized load trace of each organ, peaks were detected on the smoothed signal (0.5 min moving-mean window) using a minimum inter-peak distance of 1 min and a minimum peak prominence of 0.05 (normalized units). Interpeak intervals were computed as the time between consecutive detected peaks. Peaks were validated by visual inspection.

##### Gastric emptying classification

To classify stomach clips by whether active gastric emptying was occurring, the corresponding duodenal load trace was examined within each clip window. Clips in which duodenal load remained below 0.15 (normalized units) throughout were classified as non-emptying; clips in which the duodenal derivative exceeded 20% of the global maximum derivative were classified as emptying; clips meeting neither criterion were classified as undecided and excluded from further analysis. Antral contraction period, calculated by autocorrelation as described below, was compared between emptying and non-emptying clips.

##### Alignment and correlation with photometry

For simultaneous photometry-fluoroscopy sessions, fluoroscopy and photometry recordings were aligned based on the acquisition start times. Pearson correlation between organ load and photometry z-score was computed at a lag of 22 s, which is the average distance between duodenal and photometry peaks. Significance was assessed by comparison to a null distribution generated from 1,000 iterations of circularly shifted photometry traces. To determine NPFF neuron activity aligned to duodenal filling events, peaks in the duodenal load trace were detected (minimum prominence 0.10, minimum inter-peak distance 10 s, following additional 15 s smoothing), and photometry activity was averaged in a window surrounding each detected peak.

### Reanalysis of NTS Vglut2 single-cell calcium dynamics

Calcium imaging data from NTS glutamatergic neurons, previously collected in the lab using single-cell imaging^41^, were reanalyzed here using a new clustering approach. For each experiment, activity traces for individual neurons were extracted for each mouse, and all responses were normalized using the function z = (C_raw_ - μ)/σ, where C_raw_ is an output of the Inscopix software, μ is the mean C_raw_ during the 10 min baseline period before stimulus presentation and σ is the standard deviation of C_raw_ during the same baseline period. We defined a neuron as “activated” if the mean z-score was ≥ 1z or its 95^th^ percentile z-score > 2z. Neurons with a mean z-score of < −1z were defined as “inhibited” and a mean z-score between - 1z and 1z was defined as “non-responsive”.

Activated cells were then subdivided by activity pattern. Principal component analysis was performed on the smoothed traces of activated cells (top 20 components), and k-means clustering (50 replicates, squared Euclidean distance) was applied to the resulting PC scores; cluster number (k = 3) was selected using the elbow method on total within-cluster inertia. This yielded three clusters, corresponding to Early, Late, and Continuous response patterns, respectively.

Within the Continuous cluster, cells were further subdivided using two amplitude-normalized features computed over the post-infusion window: a trend axis, defined as the normalized difference between late- and early-window mean activity, and a peakiness axis, defined as the normalized standard deviation of the residual signal after subtracting a slow moving-median trend. K-means clustering (k = 3, fixed) was applied to these two z-scored features, yielding three subclusters that were assigned biological labels according to their relative feature values: the subcluster with the highest peakiness was labeled “pulsatile,” and the two remaining subclusters were labeled “ramping up” and “ramping down” according to whether their trend value was most positive or most negative, respectively.

### Reanalysis of MERFISH Data from the Allen Brain Atlas

Allen Brain Cell Atlas MERFISH dataset^50^ (C57BL6J-638850, primary adult mouse brain, 500-gene panel) was analyzed using the atlas’s existing whole-mouse-brain taxonomy (CCN20230722). Analysis was restricted to coronal sections containing the dorsal vagal complex (sections 5–12). The midline was set at 5.7 mm (half of the CCF’s 11.4 mm width), and each cell’s mediolateral distance from the midline was calculated as |z_ccf − 5.7|. Glutamatergic clusters containing at least 15 cells were ranked from most medial to most lateral based on the median mediolateral distance of their cells. The most medial cluster was 4546 NTS Dbh Glut_3, a noradrenergic population (A2/C2) marked by Dbh and Th expression.

To identify a gene that specifically marks this cluster, we used a companion single-cell RNA-sequencing reference dataset with the same cell-type taxonomy (CCN20230722 annotation) as our spatial data. This reference includes the average expression level of all 32,285 genes sequenced (expressed as normalized log2 values) for every cluster. We then compared cluster 4546 to the 65 other neuronal clusters in the dorsal vagal complex (DVC) that also had at least 15 cells. For each gene, we calculated a specificity score: expression in cluster 4546 − expression in the next-highest-expressing DVC cluster. Genes were ranked by this score, keeping only genes that were reasonably well-detected in cluster 4546 (log2 expression ≥ 4). The top-ranked gene was Npff: it averaged 11.6 (log2) in cluster 4546, compared to only 2.1 in the next-highest DVC cluster — the largest margin of any of the 65 clusters compared.

To confirm the spatial distribution of Npff, we used the Allen imputed MERFISH matrix, which estimates the expression of other marker genes for every cell in the whole-brain MERFISH dataset (Npff was not included in the 500-gene MERFISH panel). Consistent with the RNA-seq result, imputed Npff expression was highest in cluster 4546 (median log2 11.8, versus 0.2 across the rest of the DVC), and the cells with the highest Npff levels were in the medial DVC, the same location as cluster 4546.

### Fiber photometry recordings

#### Photometry setup

Mice were connected to a patch cable (Doric Lenses, MFP_400/460/900-0.48_2m_FCM-MF2.5) for photometry recordings. A blue LED (470 nm) and a UV LED (405 nm) served as excitation sources for the calcium-dependent GCaMP signal and the isosbestic control signal, respectively. Both LEDs were driven by a multichannel hub (Thorlabs) and sinusoidally modulated at 305 Hz and 505 Hz before being delivered through a filtered minicube (Doric Lenses, FMC6_AE(400-410)_E1(450-490)_F1(500-540)_E2(550-580)_F2(600-680)_S) and through the implanted optic fiber. Emitted fluorescence was collected back through the same fiber, separated at the dichroic ports of the minicube, and detected by a femtowatt silicon photoreceiver (Newport, 2151). The resulting signals were digitally sampled at 1.0173 kHz, demodulated by lock-in amplification, and acquired through a real-time processor (RZ5P, Tucker-Davis Technologies) using Synapse software (TDT). Data was exported using TDT Browser and downsampled to 4 Hz in MATLAB prior to analysis.

#### Behavior

For all recordings, mice were placed in isolated behavioral chambers (Med Associates) without water or food access unless otherwise specified. Chambers were cleaned between experiments with 70% ethanol. Mice were habituated for one night in the chambers before experiments with lickometers containing water and chow in the chamber. Mice received an infusion of water, and an infusion of Ensure for habituation, several days prior to the first experiment. Before each recording, photometry implants on individual mice were cleaned with 70% ethanol using connector cleaning sticks (MCC-S25) and connected to a photometry patch cable immediately afterwards. For all photometry experiments, mice were acclimated to the behavioral chamber for at least 20 min with recording before presentation of a stimulus.

Mice were fasted overnight before IG or ID infusion experiments. The IG or ID catheter was attached to a syringe pump (Harvard Apparatus, 70-2001) using plastic tubing and adapters (AAD04119, Tygon; LS20, Instech). For the nutrient panel experiments, mice received a 1 mL infusion of saline, 24% glucose, 10% intralipid (Sigma, I141-100ML; Medline, BHL2B6064H), 45% protein (Proteinex, Cat. 54859-525-30), or 24% mannitol over 10 min (0.1 mL/min). For the intermittent bolus infusion paradigm, mice received six 0.1 mL boluses of the same macronutrients, delivered every 10 min over one hour, by either the IG or ID route.

##### Intermittent lick access

To measure NPFF neuron responses during voluntary ingestion, mice were tested in the same behavioral chambers using a paradigm that matched the intermittent bolus infusion experiments. Mice were fasted overnight and acclimated to the chamber for at least 20 min before the first access period. Mice were then given 1 min of access to a lick spout containing either saline or 10% intralipid (Sigma, I141-100ML; Medline, BHL2B6064H), repeated every 10 min for a total of six access periods over one hour. Access was controlled by a retractable sipper. Licks were monitored using a contact lickometer interfaced with Med Associates Med-PC software and recorded on a channel synchronized to the photometry signal.

#### Analysis

Calcium signals were converted to %ΔF/F by fitting the isosbestic (405 nm) channel to the GCaMP (470 nm) channel by linear regression over the baseline period and normalizing to this fit. Normalized ΔF/F traces were then converted to a z-score using the mean and standard deviation of the pre-infusion baseline period. For display purposes, mean traces were downsampled by a factor of 20 or 120 to reduce file size. Baseline activity was calculated from the 10 minutes preceding stimulus presentation, during which animals remained undisturbed in the behavior chamber. Photometry interpeak intervals were computed by smoothing each animal’s z-scored trace (30 s moving mean) and detecting peaks with a minimum inter-peak distance of 60 s and a minimum prominence of 0.5. Interpeak intervals were calculated as the time between consecutive detected peaks. The mean response was calculated by comparing the average z-score during the baseline to the average z-score during the post-stimulus epoch of interest. Mean z-scores reported are for the first 10 minutes of this epoch, unless otherwise noted.

For the intermittent infusion experiments, calcium signals were converted to normalized ΔF/Ftraces and then to z-scores as described above. After, each of the si× 10-min spaced boluses per animal were extracted by a window of 2 min before to 8 min after infusion onset. Each bolus window was rebaselined independently, using the mean and SD of that bolus’s own 2-min pre-infusion period, yielding a z-score specific to that bolus. Per-animal traces were generated by averaging across all valid boluses within each animal and PSTHs were made by averaging all boluses across all animals. For each bolus, the mean z-score was calculated over the 0–2 min post-infusion window; these values were averaged across boluses within an animal, then across animals. Response time was defined as the first point at which the smoothed trace crossed 2 standard deviations above baseline and remained there for at least 1 second.

### Optogenetic experiments

#### Laser parameters

For continuous and pre-stimulation protocols, light was delivered at 20 Hz using 10-ms pulses in a 2-second on, 3-second off pattern. For closed-loop experiments during ingestion, a 2-second train of light was triggered by each detected lick, monitored using a contact lickometer interfaced with Med Associates Med-PC software. Light was delivered through a single fiber optic patch cord (Doric, MFC_200/240-0.22_6mm_ZF1.25(G)_FLT) connected to a 473-nm DPSS laser (Shanghai Laser and Optics Century BL473-100FC). Laser power was set to 10 mW, measured at the tip of the patch cord prior to each experimental session.

#### Fos mapping

Npff-Cre/+::ChR2/+ mice and controls (Cre+ChR2- or Cre-ChR2+) received continuous optogenetic stimulation for 10 min and were perfused 90 min after stimulation onset for Fos immunostaining. Two days prior to the experiment, mice received an intraperitoneal injection of Fluorogold (Fluorochrome, 20 mg/kg) to retrogradely label peripherally projecting neurons.

#### Behavior

All stimulation experiments were counterbalanced for order and included within-animal (laser) and genotype controls except for the terminal intestinal transit assay which included only genotype controls. Genotype controls were littermates lacking the Cre or reporter allele.

##### Optogenetic gastric emptying assay

Mice received an IG infusion of 0.5 mL barium/water mixture over 5 min. 30 s videos were acquired at t = 0, 5, 10, 15, 20, 25, 30, and 40 min.

##### Intestinal transit assay

Mice received an IG infusion of 0.2 mL barium/water mixture with red food dye over 1 min. 19 minutes after the start of the infusion, 2min X-ray fluoroscopy video was recorded, bracketing laser onset at minute 20 for further analysis. Laser stimulation continued for an additional 20min (40 min total from the start of infusion), after which animals were perfused and intestines dissected for measurement of dye front distance. The small intestine was excised, and the distance traveled by the dye front (measured to its furthest visible point) and the total length of the small intestine (measured from the pyloric sphincter to the cecum) were both determined using a ruler. Dye travel distance was expressed as a percentage of total small intestine length.

##### Acetaminophen absorption assay

Mice were fasted overnight prior to oral gavage of 0.5mL of 20 mg/mL acetaminophen in saline. Animals received continuous optogenetic stimulation (as described above) for the 10 min between gavage and blood collection, or no stimulation (laser off), in control and ChR2-expressing mice. 10 minutes after gavage, a single blood sample was collected via tail vein. Blood was spun at 10,000 RCF at 4°C for 10 min, and plasma was collected and frozen at −80°C. Plasma acetaminophen concentration was measured using an Acetaminophen L3K detection kit (Sekure Chemistry) on a Multimode Microplate Reader (Agilent BioTek Synergy LX) and reported as mg/dL.

##### Body temperature assay

Mice were implanted subcutaneously in the dorsal flank with a temperature transponder (UCT-2112, UID) using a pistol grip injector (UPGI-Q, UID) under anesthesia at least one week prior to experiments. During optogenetic stimulation sessions, body temperature was recorded using a handheld reader (URH-2, UID) every 5–10 minutes over an 80-minute session, with continuous laser stimulation delivered from minute 10 to minute 70.

##### Feeding assays

Mice were tested in behavioral chambers (Med Associates) without food or water access, and chambers were cleaned with 70% ethanol between sessions. Before experiments began, mice underwent two habituation sessions: an overnight session with ad libitum access to water and chow, followed the next day by a second session in which animals were connected to the fiber optic patch cord for an hour with the laser off. For Ensure, glucose (24%), and intralipid (10%) licking experiments, as well as for chow and HFD experiments, mice were given access to each compound for at least 24 hours in their home cage a few days prior to the experiment to prevent neophobia.

For chow and HFD consumption experiments, animals were fasted overnight before the experiment. After a 10-min acclimation period in the chamber, animals received a pellet of standard chow (PicoLab 5053) or HFD (Research Diets, D12492) for self-paced consumption over 1 hour, with continuous optogenetic stimulation delivered throughout the session or no laser treatment.

For glucose and intralipid licking experiments, mice were fasted overnight prior to testing. On the experimental day, animals were given a 10-minute acclimation period in the chamber before liquid access and stimulation. Liquid access and stimulation then occurred simultaneously for 1 hour.

For closed-loop stimulation, animals were given 60 min of access to Ensure in which a laser pulse train was triggered by each detected lick, such that stimulation occurred only during licking. For pre-stimulation, mice received 30 min of continuous optogenetic stimulation in the behavioral chamber prior to food access. Following stimulation, animals were given 60 min of access to Ensure without further light delivery.

##### Cholinergic blockers

Mice were fasted overnight and received an intraperitoneal injection of vehicle (10% DMSO, 10% Tween-80 in saline), methylatropine bromide, 10mg/kg (Millipore Sigma) or darifenacin hydrobromide, 5mg/kg (Millipore Sigma). 5 minutes after injection, animals received continuous optogenetic stimulation or no stimulation (laser on vs. off) and were given access to Ensure for 60 min.

#### Analysis

##### Gastric emptying assay

Stomach area was quantified using a custom MATLAB tool. Briefly, a calibration frame containing a ruler of known length was used to establish a pixel-to-millimeter conversion factor, and a time-zero frame corresponding to the start of infusion or optogenetic stimulation was designated for each video. For each analyzed frame, a freehand region of interest was drawn around the stomach; within this region, the image was Gaussian blurred, normalized, and thresholded using Otsu’s method to generate candidate dark and bright regions, with the lower-intensity region selected as the stomach mask. Holes were filled and small objects removed, the largest connected component was retained, and its area was computed in square millimeters using the calibration factor. Each segmentation was visually inspected and accepted or repeated until satisfactory.

Stomach area was measured at the multiple time points described above for each animal. Antral contraction period was extracted from the raw mask area trace by autocorrelation. For this assay, a clip beginning at minute 10 after the start of infusion was visually reviewed for each animal and condition to identify sessions in which antral anatomy permitted visualization of contractions, and clips meeting this criterion were segmented using SAM2 and analyzed by the same method used for fluoroscopy above.

##### Intestinal Transit Assay

Organ load for this assay was computed as described above, with load traces normalized to the pre-laser baseline period rather than to session maximum, yielding the change-from-baseline values.

##### Feeding behavior

Lick microstructure was analyzed using a custom Matlab pipeline. A licking bout was defined as a continuous sequence of at least 3 licks in which no inter-lick interval exceeded 5 seconds. Total lick count, latency to first lick, bout number, and mean bout duration were computed for each animal across the full session. Cumulative lick counts were computed in 10-second bins aligned to the time of lickometer access and averaged across animals within each group. To measure chow and HFD consumption, the pellet was weighed before and after the experiment.

### EMG recordings

#### EMG setup

All recordings were performed in urethane-anesthetized mice (*Npff-Cre::Ai32* or littermate controls). Mice received an oral gavage of 10% intralipid 5 minutes before anesthesia was induced via intraperitoneal (i.p.) injection of urethane (1.5 g/kg) and placed on a heating pad. Anesthesia was confirmed by the absence of withdrawal reflexes to toe and tail pinch prior to surgery.

The abdominal cavity was exposed, and the sternum was lifted using a sternum clamp to improve visual access of the diaphragm and upper gastrointestinal tract. Two additional bilateral retractors were used to elevate the sides of the abdominal wall, allowing the gastrointestinal tract to remain covered with saline throughout the experiment.

Twelve single-ended electrodes were used for recordings. Two electrodes were placed in the diaphragm muscle to capture cardiac (electrocardiogram, ECG) and diaphragmatic (breathing) activity, and ten additional electrodes were placed along the gastrointestinal tract (3 in stomach, 5 in proximal duodenum, and 2 or 3 in distal small intestine), positioned using anatomical landmarks (e.g. pylorus). All recording electrodes were referenced to a single reference electrode secured between the skin and lower ribcage, with a subcutaneous ground electrode placed near the thigh. The diaphragm and ECG recordings were used during offline analysis to identify and remove motion-related artifacts from gastrointestinal channels.

Recording electrodes were made of Teflon-coated tungsten wire (A-M Systems, #795500). Teflon insulation was removed from the wire tip (<1.0 mm), and the exposed tip was formed into a hook and passed through the target tissue. Electrodes were connected to a 36-pin wire adapter (RHD Part #C3420) attached to a 64-channel RHD headstage (Intan, #C3315). Data were acquired at 30 kHz using an RHD 512-channel Recording Controller (Intan Technologies) and RHX Software (version 3.1.0), with a fixed hardware gain of 192. A 60 Hz notch filter was applied online during acquisition.

Following a 5-minute baseline recording, optogenetic stimulation was delivered using a 473-nm laser. Stimulation consisted of 10-ms pulses delivered at 20 Hz in a 2-second on, 3-second off pattern for 2 minutes. Laser power was set to 10 mW, measured at the tip of the patch cord prior to each experimental session. Laser pulses were triggered by TTL pulses generated through an Arduino and were simultaneously acquired through a synchronized digital input channel.

#### Analysis

Each recording channel was truncated to a fixed 15-minute period (5 minutes before laser onset and 10 minutes after laser onset) prior to analysis. Raw int16 traces were converted to microvolts using the manufacturer specified gain (0.195μV/bit). All recordings were aligned to the first rising edge of the laser TTL channel. Analysis was then performed in three consecutive two-minute windows: two minutes immediately preceding laser-onset, a two-minute laser stimulation period spanning the full stimulation epoch, and a two-minute window immediately following the end of laser stimulation.

Non-rhythmic, large amplitude ECG channel events consistent with gasping or motion artifact were identified separately from cardiac beats prior to gut spike event detection. If a gasping event was detected, a 1 second window around the event was removed across all channels for that animal to avoid accidental spike potential detection. Across all mice, gasp-related exclusion accounted for less than 1% of the analyzed recording period (maximum of 3.6 s omitted from a 6-minute analysis window per animal).

Prior to event detection, EMG channels were screened for baseline instability. For each recording channel, the pre-stimulation baseline (before laser onset) was assessed by calculating the moving root mean square (RMS) amplitude using a 30-s sliding window. A linear regression was fitted to the baseline RMS trace, and the percentage change in RMS across the baseline period and the coefficient of determination (R²) were calculated. Channels exhibiting both a large positive baseline drift (greater than the cohort median plus three median absolute deviations) and a linear trend (R² > 0.3) were automatically excluded. Additional channels with visually confirmed recording instability were removed by manual inspection before downstream analyses.

Spike potentials were detected independently for each gut EMG channel, and applied identically across all channels and groups, with gasp windows from above excluded from the detection input. For each channel, the cleaned signal was band-pass filtered at 10-200 Hz using a 4^th^ order Butterworth filter. Cardiac artifact was blanked by linear interpolation across a 10 ms window around each detected heartbeat. A per-channel detection threshold was set at 5 times the standard deviation of the filtered baseline period. Contiguous samples exceeding threshold were grouped into candidate spike potentials. Since most if not all spike potentials exceeded 20 ms, spikes less than 6 ms were rejected as residual ECG breakthrough. Events occurring within 25 ms of one another were merged, retaining the larger-amplitude detection to avoid double counting the two phases of a single spike potential. This threshold-per-channel approach was chosen to favor undercounting over overcounting of ambiguous low-amplitude events. Detected spike times, amplitudes, and polarities were retained for all channels, including channels with no significant activity, to support downstream quality control. Spike potential rates are reported as counts per minute, normalized within each channel’s respective 2-min window.

Heart rate was computed for each animal from its diaphragm ECG channel by detecting R-peaks and was expressed in beats per minute. Heart rate was calculated in the same three 2-min windows used for spike analysis (pre-stimulation, stimulation, and post-stimulation). Within the stimulation window, heart rate was additionally computed separately for the laser-ON (2 s) and laser-OFF (3 s) sub-epochs, which alternated 24 times across the 2-min stimulation period.

### Chemogenetic experiments

#### Behavior

All experiments were counterbalanced for order and included within-animal (vehicle vs. DCZ) and genotype (±DREADD) controls, with genotype controls being littermates lacking the Cre or reporter allele. Animals were tested in behavioral chambers (Med Associates) without food or water access, and chambers were cleaned with 70% ethanol between sessions. Before experiments began, mice underwent an overnight habituation session with ad libitum access to water and chow followed the next day by a second session in which animals received a saline injection and then licked Ensure in the chamber.

For Ensure, glucose (24%), and intralipid (10%) licking experiments, as well as for chow and HFD experiments, mice were given access to each compound for at least 24 hours in their home cage a few days prior to the experiment to prevent neophobia. Mice were fasted overnight prior to the experiment. On the experimental day, mice were injected intraperitoneally with either deschloroclozapine (DCZ, 0.3 mg/kg, Hello Bio) or saline vehicle, then placed in the behavioral chamber for a 10-min acclimation period before food presentation.

#### Analysis

Licking bout analysis was performed as described above. To measure chow and HFD consumption, the pellet was weighed before and after the experiment.

### Histology

Mice were transcardially perfused with heparinized PBS followed by 10% formalin. Brains were post-fixed overnight in 10% formalin at 4°C, then transferred to 30% sucrose in PBS overnight. Brains were sectioned at 30 μm (for vagal afferent visualization), 40 μm (for IHC) or 15μm (for ISH) on a cryostat and mounted with DAPI Fluoromount-G (Southern Biotech), then imaged using the Nikon Eclipse Ti2-E. Sections to visualize vagal afferents were imaged using a LSM 510 confocal.

#### In situ hybridization

In situ hybridization was performed using RNAscope Fluorescent Multiplex Kit (Advanced Cell Diagnostics, 320850) following the manufacturer’s instructions. Briefly, sections were fixed in 4% paraformaldehyde (15 min at 4°C), washed in PBS, dehydrated in a series of ethanol washes, and then dried. A hydrophobic barrier was drawn around the section with an ImmEdge pen (Vector Lab, H-4000). The sections were treated with Protease IV in a HybEZ Humidity Control Tray (30 min at RT), incubated with target probes (Mm-Npff, 479901; tdTomato-C2, 317041-C2) in a HybEZ Oven (2 h at 40°C), and then treated with Hybridize Amp 1–4. Sections were stained using TSA Vivid Fluorophore Dyes from the manufacturer (Advanced Cell Diagnostics; 32327[1/2/3]).

#### Npff^cre^ expression mapping

*Whole-brain quantification of tdTomato expression (QUINT):* The spatial distribution of tdTomato+ Npff neurons (Npff^cre^::Ai14) was quantified from serial coronal sections using the QUINT workflow. Section images were registered to the Allen Mouse Brain Atlas (CCFv3, 2017) in a two-step procedure: sections were first manually aligned to the reference atlas using QuickNII, followed by non-linear, in-plane refinement using VisuAlign. Registered images were then segmented in Ilastik in two stages: pixel classification to distinguish tdTomato+ signal from background, followed by object classification to separate labeled cells from artifacts.

Because Npff neurons are concentrated at the interface between the AP and the NTS (most medial part of the caudal NTS), we defined an anatomical “subpostrema” border zone to add to the Allen taxonomy. Binary masks of the AP (Allen structure 207) and NTS (structure 651) were extracted, and a three-dimensional Euclidean distance transform was computed from each mask. The subpostrema was defined as the zone extending 50 µm into the AP and 150 µm into the NTS from their shared boundary. This zone was carved out of both parent regions, and cells were classified as AP core, NTS core, subpostrema, or other.

Registered and segmented images were combined and imported into Nutil Quantifier for automated quantification of tdTomato+ pixel density across all brain regions. Load was defined as the number of segmented object pixels divided by total region pixels for each atlas region (Nutil convention), with anatomical subregions collapsed to their parent structures using a pixel-weighted mean. Load for the subpostrema was computed from object and region pixels falling within the border zone, and the corresponding pixels were removed from the AP and NTS core values. To identify regions with enriched Npff labeling without an arbitrary percentile cutoff, each region’s Load was expressed relative to whole-brain background labeling density (total object pixels divided by total region pixels across all regions, 0.0011). Regions exceeding three times background (Load ≥ 0.0033) were considered enriched, identifying 15 enriched regions, with the subpostrema most strongly labeled (94× background).

#### Immunohistochemistry

Floating sections were washed three times in PBS-T (0.1% Triton-X), blocked in 2% normal donkey serum in PBS-T for one hour at room temperature, and incubated overnight in primary antibodies in blocking solution at 4°C (rabbit anti-cFos, Abcam ab190289, 1:1000; chicken anti-GFP, Aves Labs GFP79749381, 1:3000; chicken anti-GFP, abcam ab13970, 1:2000). Sections were then washed in PBS-T three times and incubated in secondary antibodies (donkey anti-rabbit Alexa Fluor 568, Invitrogen A10042, 1:1000; donkey anti-chicken Alexa Fluor-488, 1:1000, Sigma-Aldrich SAB4600031) in blocking solution for one hour at room temperature, washed in PBS three times, mounted onto slides, and imaged.

Fos+ cell density in the DMV, PBN, subpostrema, rNTS, and cNTS, and the percentage of Fluorogold+ cells that were also Fos+ in the DMV, were quantified manually using the Cell Counter plugin in ImageJ.

### Statistics

All values are reported as mean ± SEM (error bars or shaded areas), unless stated otherwise. In figures, asterisks denote statistical significance: *p < 0.05, **p < 0.01, ***p < 0.001, ****p < 0.0001. Normality was assessed using the Shapiro-Wilk test; parametric tests were used for normally distributed data, and non-parametric tests were used otherwise. For paired or unpaired two-group comparisons, p values were calculated using paired or unpaired t-tests (parametric) or the Wilcoxon signed-rank test or Mann-Whitney U test (non-parametric), as appropriate.

For comparisons across multiple groups, p values were calculated using two-way ANOVA with Šidák’s multiple comparisons test or Dunnet’s multiple comparisons test, or the Kruskal-Wallis test with Dunn’s multiple comparisons test, as appropriate. To compare two distributions, the Kolmogorov-Smirnov test was used, and the Bonferroni correction was used to correct for multiple comparisons. Correlations between photometry and organ load were assessed by Pearson correlation and tested against a null distribution generated from 1,000 circularly shifted traces. Linear relationships were assessed by simple linear regression fitted by least squares. No statistical method was used to predetermine sample size. Experiments were not randomized or blinded. Photometry, behavioral, and fluoroscopy data were analyzed in MATLAB, Python, and GraphPad Prism. The statistical test, test statistic, degrees of freedom, and n used for every panel are reported in Table S1.

## Acknowledgements

We thank members of the Knight laboratory for discussions. We thank J. Grove, A. Buckley and T.J. Aitken for comments on the manuscript. This work was supported by NIH grants R01-DK106399, R01-DK138127, and R01DK145100 (Z.A.K.) and UCSF William K. Bowes, Jr. and Ute Bowes Discovery Fellowship (N.D.). N.D. is a Boehringer Ingelheim Fonds PhD Fellow. Z.A.K. is an Investigator of the Howard Hughes Medical Institute.

## Author Contributions

N.D. and Z.A.K. conceived the project and designed experiments. N.D. led the experiments and analyzed the data. N.D. performed all fluoroscopy experiments. N.D., K.X., M.K., Q.L. and A.S. performed photometry, optogenetic and chemogenetic experiments. N.D. performed all intracranial surgeries. N.D. and L.Q. performed intragastric and intraduodenal catheter surgeries. N.D., I.A. and K.Y. developed and performed gut EMG experiments. B.C.J., H.N., A.L.C., and A.S.R. provided guidance on fluoroscopy, gastrointestinal assays, and data analysis. N.D., M.K., B.C.J. collected and analyzed histology. N.D., T.L. and J.Y.O. characterized the Npff line. N.D. and Z.A.K. wrote the manuscript with input from all authors.

## Competing financial interests

The authors declare no competing interests.

## Additional information

Correspondence and requests for materials should be addressed to Zachary A. Knight.

## Data availability

The data from this study are available from the corresponding author on reasonable request.

## Supplementary video legends

**Video S1. Representative fluoroscopy video showing the dynamics of the stomach and intestines during digestion.** Example fluoroscopy video taken after IG infusion of a barium/intralipid mixture. Organ masks generated by SAM2 (Segment Anything Model 2) overlaid as colored outlines: stomach (green), proximal duodenum (pink), and distal small intestine (blue). Video is shown at 2x real-time speed.

**Video S2. Antral contractions during a non-emptying epoch.** Left, X-ray fluoroscopy video taken after an IG infusion of barium/intralipid with the stomach outlined in pink. Right, corresponding normalized stomach load trace (y-axis) over time in seconds (x-axis). Periodic antral contractions (∼8–10 s intervals) are visible in both the video and the trace, without any content emptying into the proximal duodenum. Video is shown at 4x real-time speed.

**Video S3. NPFF neuron activity tracks duodenal filling in real time.** Left, X-ray fluoroscopy video taken after IG infusion of a barium/intralipid mixture. The proximal duodenum is outlined in pink. Right, corresponding traces over time (s): top, proximal duodenal load (pink), showing filling of the proximal duodenum as gastric contents empty; bottom, simultaneous NPFF photometry signal (z-scored, green). NPFF activity closely tracks duodenal filling in real time. Video is shown at 2x real-time speed.

**Video S4. Optogenetic stimulation of NPFF neurons inhibits gastric emptying.** X-ray fluoroscopy videos from the same ChR2+ animal on separate sessions, both 25 min after infusion of a barium/water mixture. Left, laser off: the stomach is smaller and actively emptying, with antral contractions driving gastric contents into the proximal duodenum. Right, laser on: the stomach remains distended and gastric emptying is minimal while antral contractions continue to happen. Video is shown at 2x real-time speed.

**Video S5. Optogenetic stimulation of NPFF neurons drives proximal duodenum emptying.** X-ray fluoroscopy video from a ChR2+ animal 20 minutes after IG infusion of a water/barium mixture with arrows indicating the stomach and proximal duodenum. During the laser-off period (black text, top), duodenal contents mix in place without emptying distally. Upon laser onset (text switches to ‘Laser ON’ in blue), duodenal contents empty rapidly, showing that NPFF stimulation accelerates proximal duodenal clearance. Video is shown at 4x real-time speed.

**Video S6. Optogenetic stimulation of NPFF neurons induces contractions in the proximal duodenum.** Left, schematic of the experimental setup. Right, video of the exposed proximal duodenum (outlined by white circle) in an anesthetized animal. Laser onset is indicated by ‘Laser ON’ text appearing (blue) and flashing of blue laser in the background; duodenal contractions begin upon laser onset. Video is shown at 2x real-time speed.

---

**Supplementary Figure 1.**
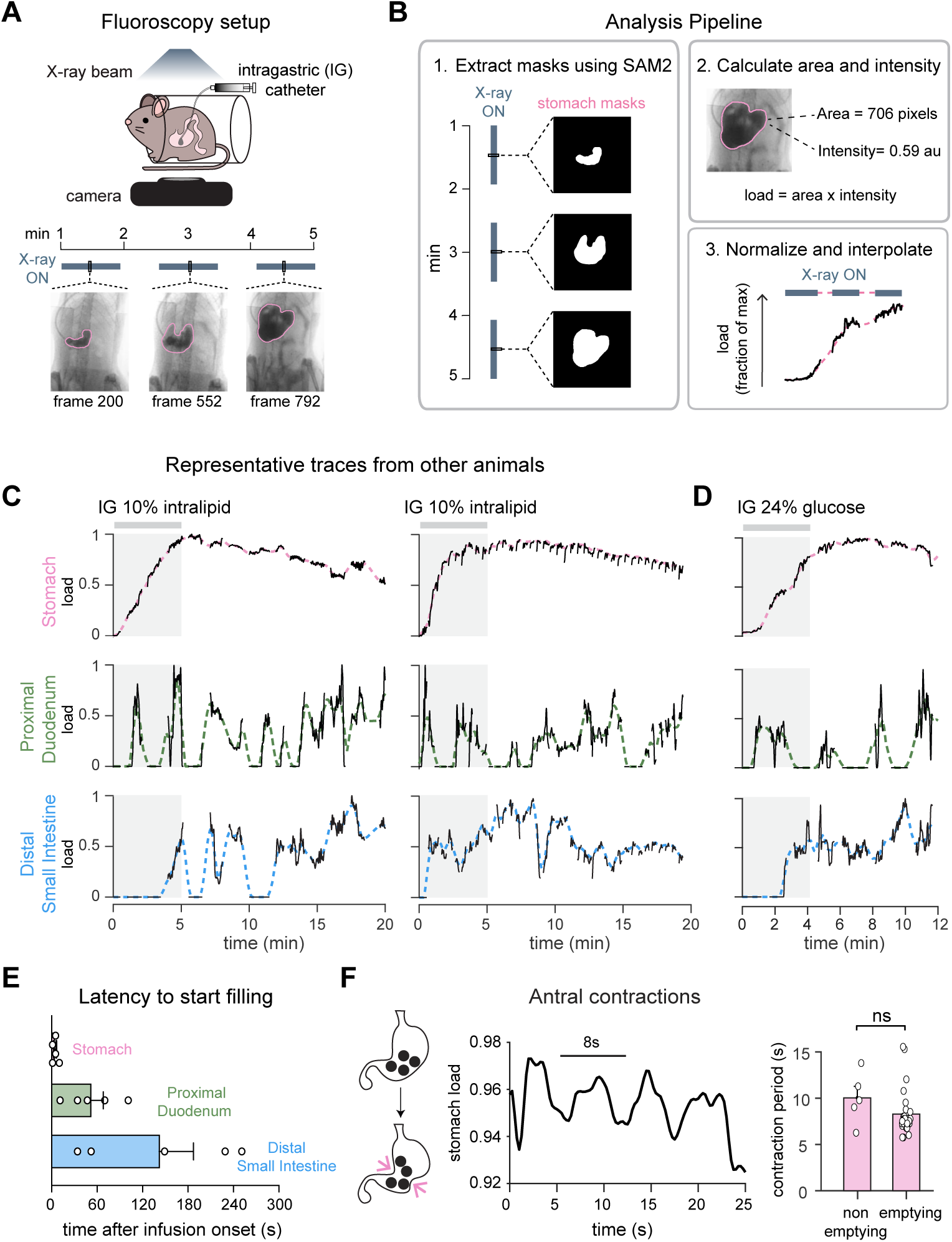
Quantification of GI transit dynamics by x-ray fluoroscopy. **A.** Fluoroscopy setup. Mice were infused intragastrically via catheter with a barium/nutrient mixture and imaged under X-ray. X-ray exposure was intermittent to limit radiation exposure. X-ray ON periods are indicated by gray bars. Bottom, representative fluoroscopy frames at three timepoints. **B.** Analysis pipeline for quantifying GI organ load from fluoroscopy videos. (1) Organ masks were extracted from each frame using SAM2. (2) Area and background-subtracted intensity within each mask were calculated and multiplied to yield load. (3) Load traces were interpolated across discontinuous clips and normalized to fraction of max (see methods). **C.** Representative normalized load traces for the stomach (pink), proximal duodenum (green), and distal small intestine (blue) from two additional mice during IG infusion of 10% Intralipid (gray bar). **D.** Representative normalized load traces from a mouse infused with 24% glucose (gray bar). **E.** Latency to start filling for each compartment, measured from infusion onset (n=5 mice). **F.** Antral contraction rate is not modulated by gastric emptying state. Left, schematic of antral contractions propelling stomach content into the duodenum. Middle, representative stomach load trace showing rhythmic (every ∼8-10 s) antral contractions. Right, contraction period during non-emptying and emptying epochs. Each dot represents an xray ON epoch. ns, not significant.|

**Supplementary Figure 2.**
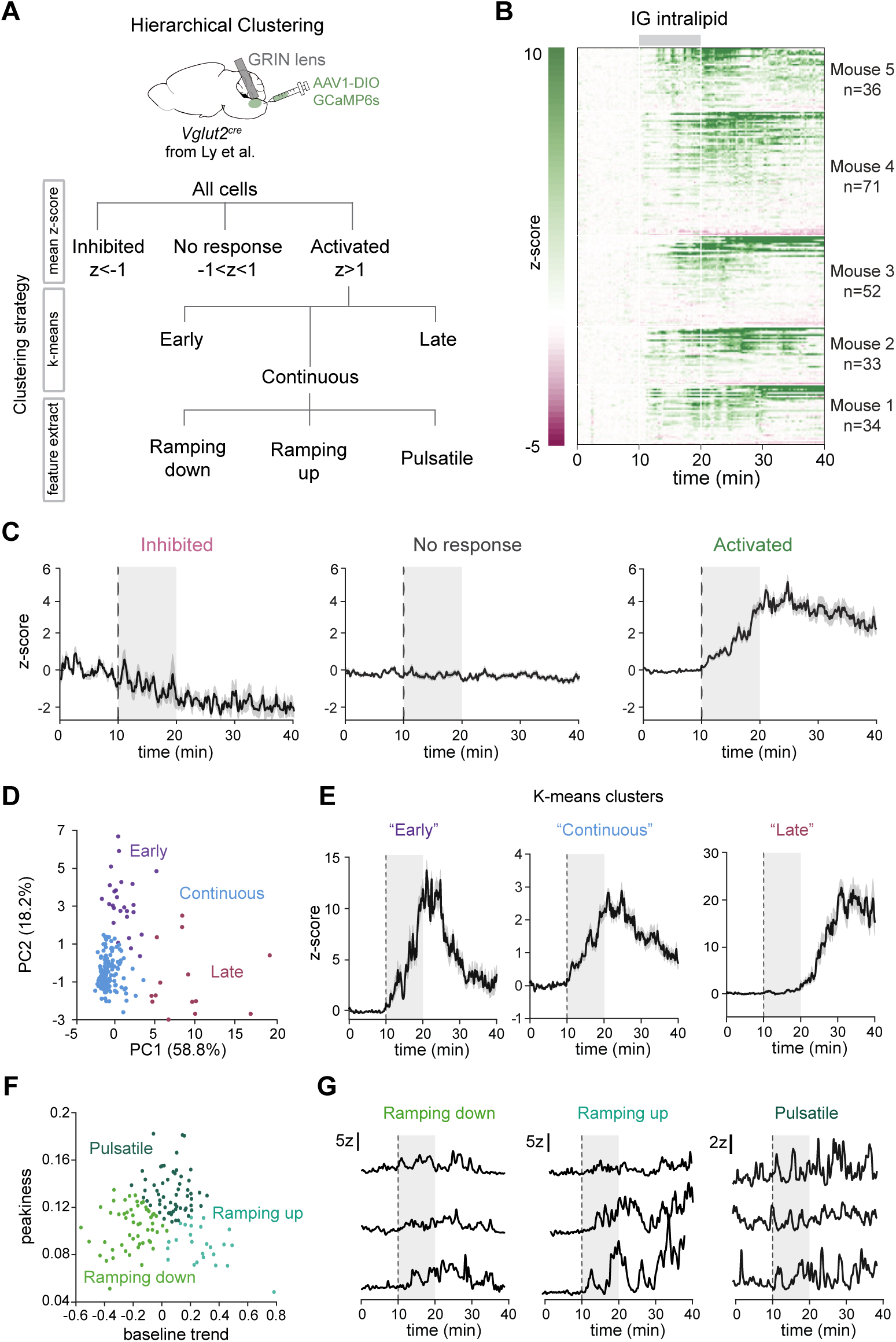
Hierarchical classification of NTS Vglut2 neuron responses to intragastric infusion of intralipid. **A.** Hierarchical clustering strategy for NTS Vglut2 neurons imaged during intragastric (IG) intralipid infusion. Top, schematic of single-cell imaging (GRIN lens, AAV1-DIO-GCaMP6s in *Vglut2^cre^* mice). Bottom, clustering tree. Cells were first sorted by mean z-score during infusion (Inhibited, No response, Activated), then the Activated group was split by k-means into Early, Continuous, and Late responders, and the Continuous group was further divided by extracted response features into Ramping down, Ramping up, and Pulsatile clusters. **B.** Heatmap of z-scored activity for all recorded neurons during IG intralipid infusion, grouped by mouse (n = 5 mice). **C.** Mean z-scored activity for the Inhibited, No response, and Activated clusters. **D.** K-means clustering of Activated neurons in PC space (PC1 and PC2 with % variance explained), colored by cluster (Early, Continuous, Late). **E.** Mean z-scored activity for the Early, Continuous, and Late k-means clusters. **F.** Feature-based clustering of Continuous neurons by baseline trend and peakiness, colored by cluster (Ramping down, Ramping up, Pulsatile). Baseline trend reflects the direction of change in each cell’s response (ramping down or ramping up), peakiness reflects the amplitude of fast fluctuations. **G.** Representative individual z-scored traces from the Ramping down, Ramping up, and Pulsatile clusters. Scale bars, 5z (Ramping down, Ramping up), 2z (Pulsatile). Data are mean ± s.e.m.

**Supplementary Figure 3.**
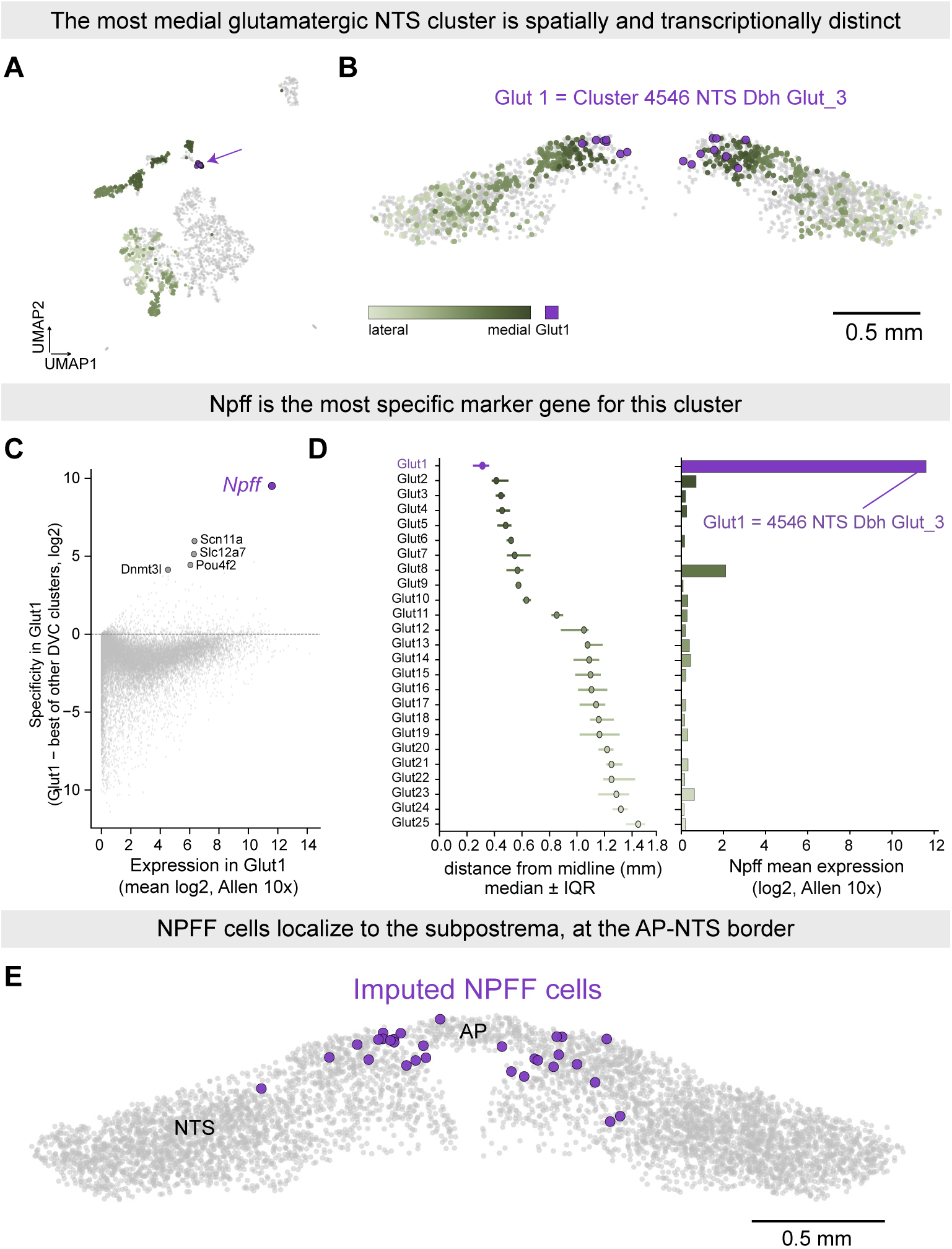
*Npff* marks a spatially and transcriptionally distinct glutamatergic cluster at the NTS-AP border. **A.** UMAP of glutamatergic neurons in the dorsal vagal complex (DVC), from the Allen Brain Cell Atlas MERFISH dataset, colored by each cluster’s median mediolateral distance from the midline (light green, lateral; dark green, medial; purple, most medial). Arrow indicates Glut1 (cluster 4546 *NTS Dbh Glut_3*), which forms a transcriptionally distinct island separate from the other clusters. **B.** Spatial distribution of DVC glutamatergic neuron populations colored as in A. Glut1 cells (purple) are concentrated at the NTS-AP border (subpostrema). **C.** Marker gene search for Glut1. For each gene, a specificity score was calculated as expression in Glut1 minus expression in the next-highest-expressing cluster. *Npff* had the highest specificity score of any gene compared. **D.** Left, distance from the midline (mm, median ± IQR) for each of 25 glutamatergic DVC clusters, ranked from most medial (Glut1) to most lateral (Glut25). Right, *Npff* mean expression (log2, Allen 10x scRNA-seq) for the same clusters in the same rank order. *Npff* expression is restricted almost exclusively to Glut1. **E.** Imputed NPFF cells (purple) overlaid on a representative DVC section. NPFF cells localize to the region spanning the area postrema (AP) and the medial NTS, consistent with the subpostrema location of cluster 4546.

**Supplementary Figure 4.**
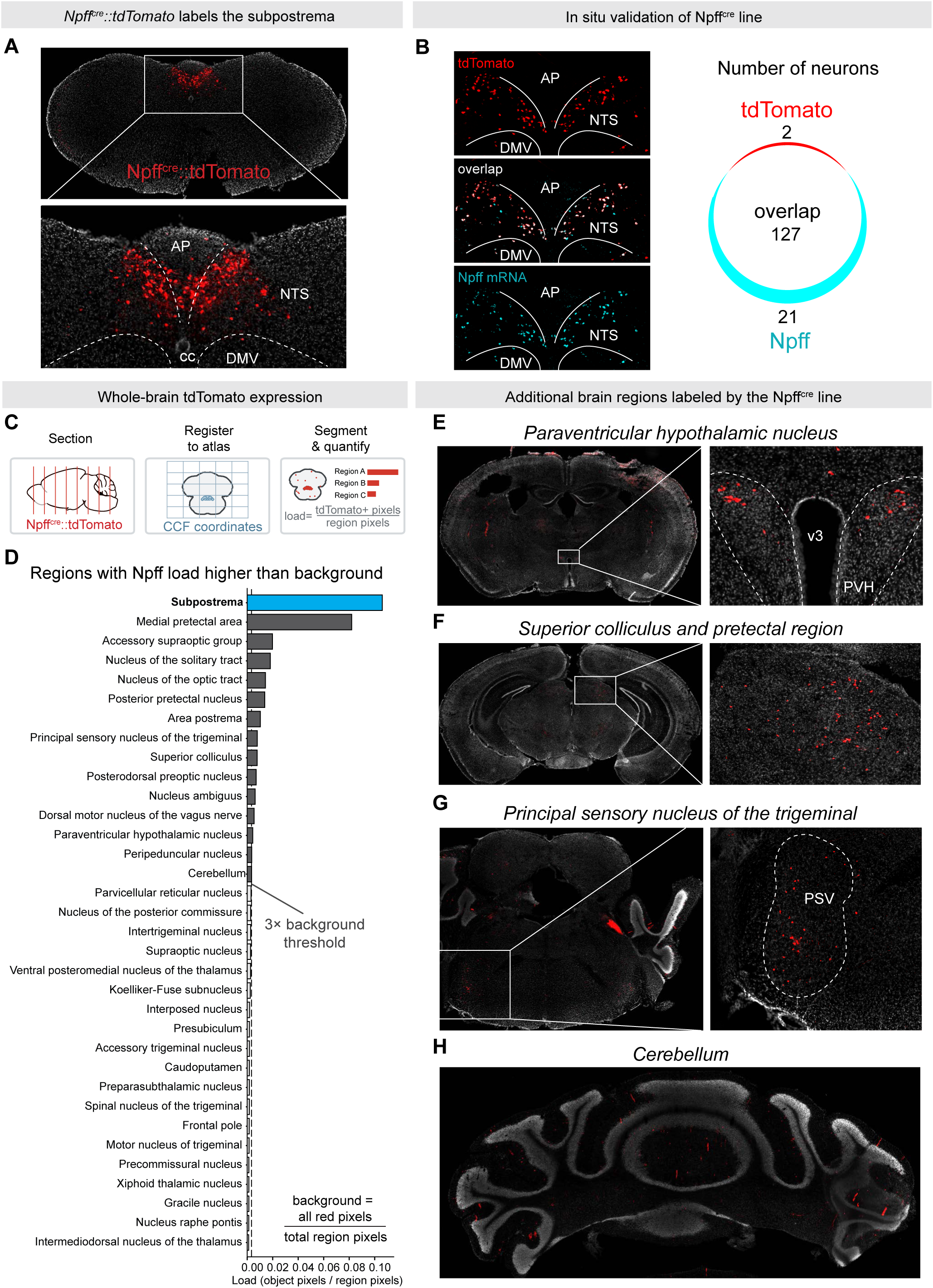
*Npff^cre^* line selectively labels subpostrema Npff neurons in the dorsal vagal complex. **A.** Representative coronal section of an *Npff^cre^::tdTomato* mouse showing tdTomato+ neurons (red) localized to the subpostrema adjacent to the area postrema (AP), nucleus of the solitary tract (NTS), and dorsal motor nucleus of the vagus (DMV). **B.** Validation of *Npff^cre^* specificity by in situ hybridization (RNAScope). Left, tdTomato (red, top) and *Npff mRNA* (cyan, bottom) signal in the same section. Overlap (white, middle) between tdTomato and *Npff mRNA* channels. Right, quantification of overlap between tdTomato+ and *Npff mRNA+* cells. Of 129 tdTomato+ cells, 98.4% co-expressed *Npff mRNA*; of 148 *Npff mRNA+* cells, 85.8% co-expressed tdTomato. **C.** Whole-brain expression of tdTomato was quantified using the QUINT pipeline: coronal sections from a *Npff^cre^::tdTomato* mouse brain were serially registered to the Allen Common Coordinate Framework (CCF) atlas, and tdTomato+ pixels were segmented and expressed as the fraction of tdTomato+ pixels relative to total region pixel (load). **D.** Ranked tdTomato load by brain region. Top region is the subpostrema. Dashed line indicates 3x background threshold, background is defined as total tdTomato + pixels divided by total brain region pixels in the dataset. Subpostrema is not a defined region in the Allen CCF; it’s designated using the combined borders of the AP and NTS (see Methods). **(E–H)** Representative images of additional brain regions in which tdTomato+ load was above the 3x background threshold in D and high expression was confirmed by visual inspection. E, Paraventricular hypothalamic nucleus (PVH), adjacent to the third ventricle (v3). F, Superior colliculus and pretectal region. Pretectal region encompasses several adjacent regions, including the medial pretectal area, posterior pretectal nucleus, and nucleus of the optic tract. G, Principal sensory nucleus of the trigeminal (PSV). H, Cerebellum.

**Supplementary Figure 5.**
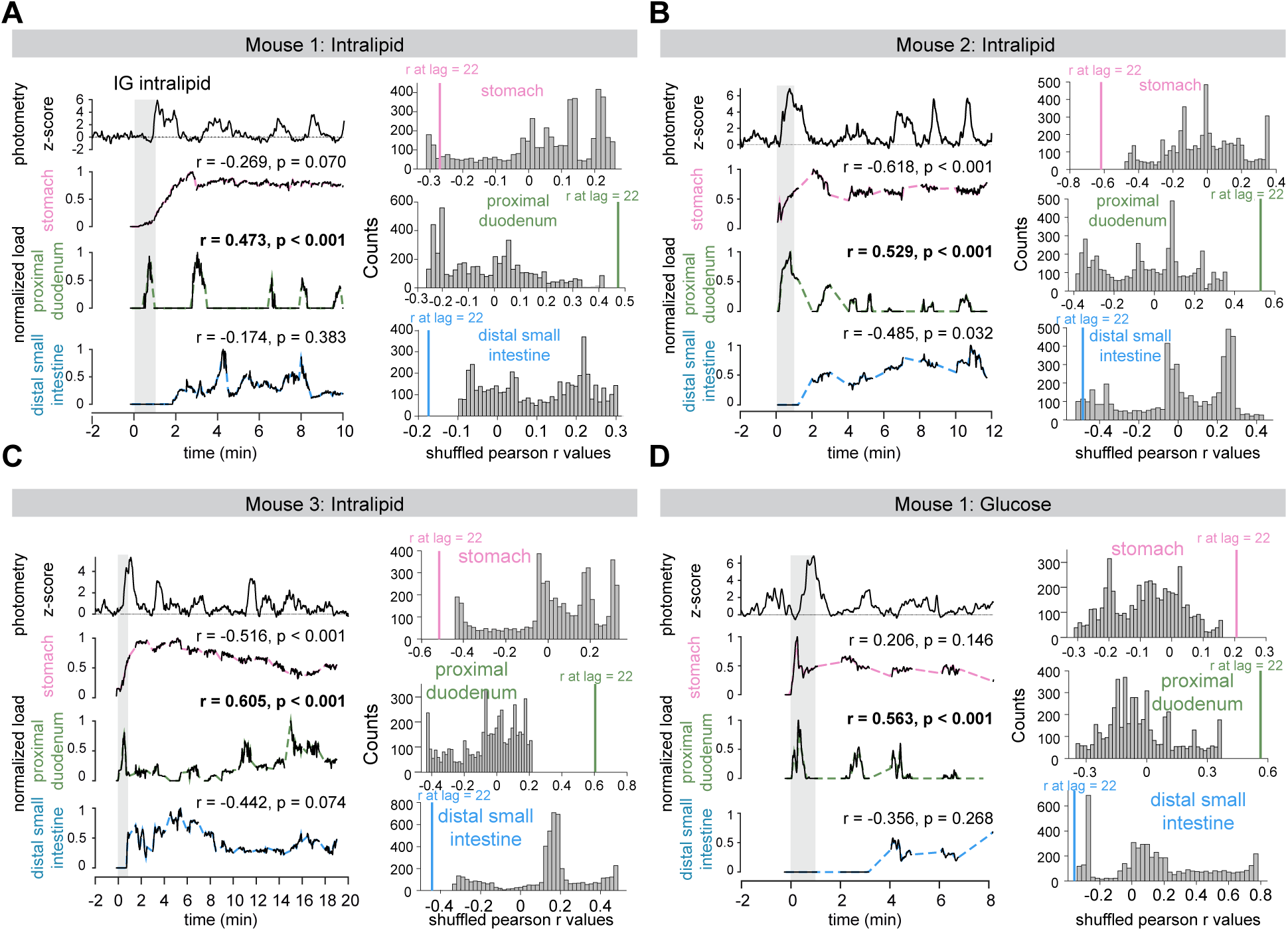
NPFF activity correlates with proximal duodenal filling across individual mice. **A–D.** Left, Npff photometry z-score trace (top) and normalized load traces for stomach (pink), proximal duodenum (green), and distal small intestine (blue, bottom) during a representative IG infusion trial in Mouse 1 (A), Mouse 2 (B), and Mouse 3 (C) infused with 10% Intralipid. Mouse 1 was also infused with 24% glucose (D). Gray shading indicates the IG infusion period. r and p values indicate the Pearson correlation between the photometry trace and each compartment’s content trace at a 22 s lag, corresponding to the average distance between NPFF and duodenal peaks (Fig. 2J). Right, distribution of Pearson r values from 1,000 iterations of circularly shuffled photometry traces for each compartment (gray). Colored vertical line indicates the true r value at the 22 s lag.

**Supplementary Figure 6.**
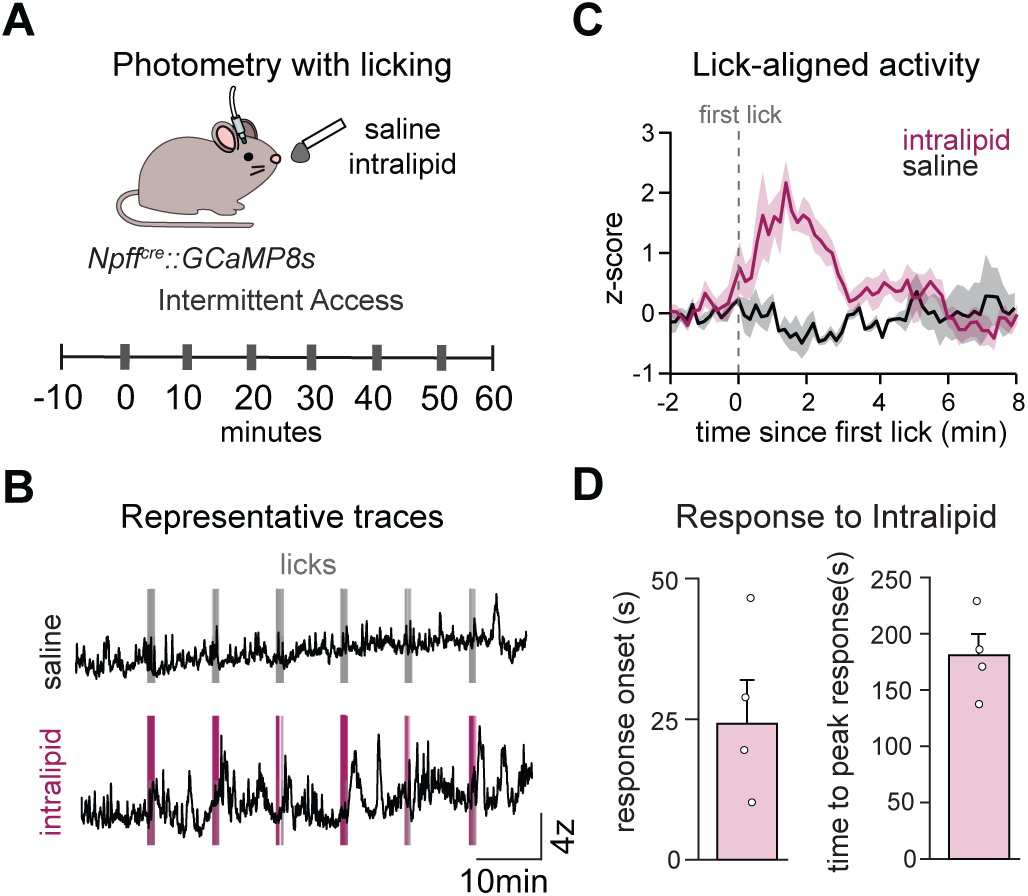
NPFF neurons are slowly activated by oral intake of intralipid, consistent with post-ingestive rather than orosensory signaling. **A.** Intermittent access paradigm. Npff^cre^::GCaMP8s mice were given 1 min access to a lick spout delivering saline (n=3) or 10% intralipid (n=4) every 10 minutes for a total of 6 times across an hour. **B.** Representative photometry traces (z-scored) from a saline session (top) and an intralipid session (bottom) from the same recording paradigm; vertical bars mark individual licks. **C.** Average NPFF response aligned to the first lick of each bout, comparing saline (black) and intralipid (pink) sessions. **D.** Quantification of the timing of intralipid-evoked response across animals: onset latency of the response (left) and time to peak response (right), relative to first lick. Each dot represents a single mouse. Data are mean ± s.e.m.

**Supplementary Figure 7.**
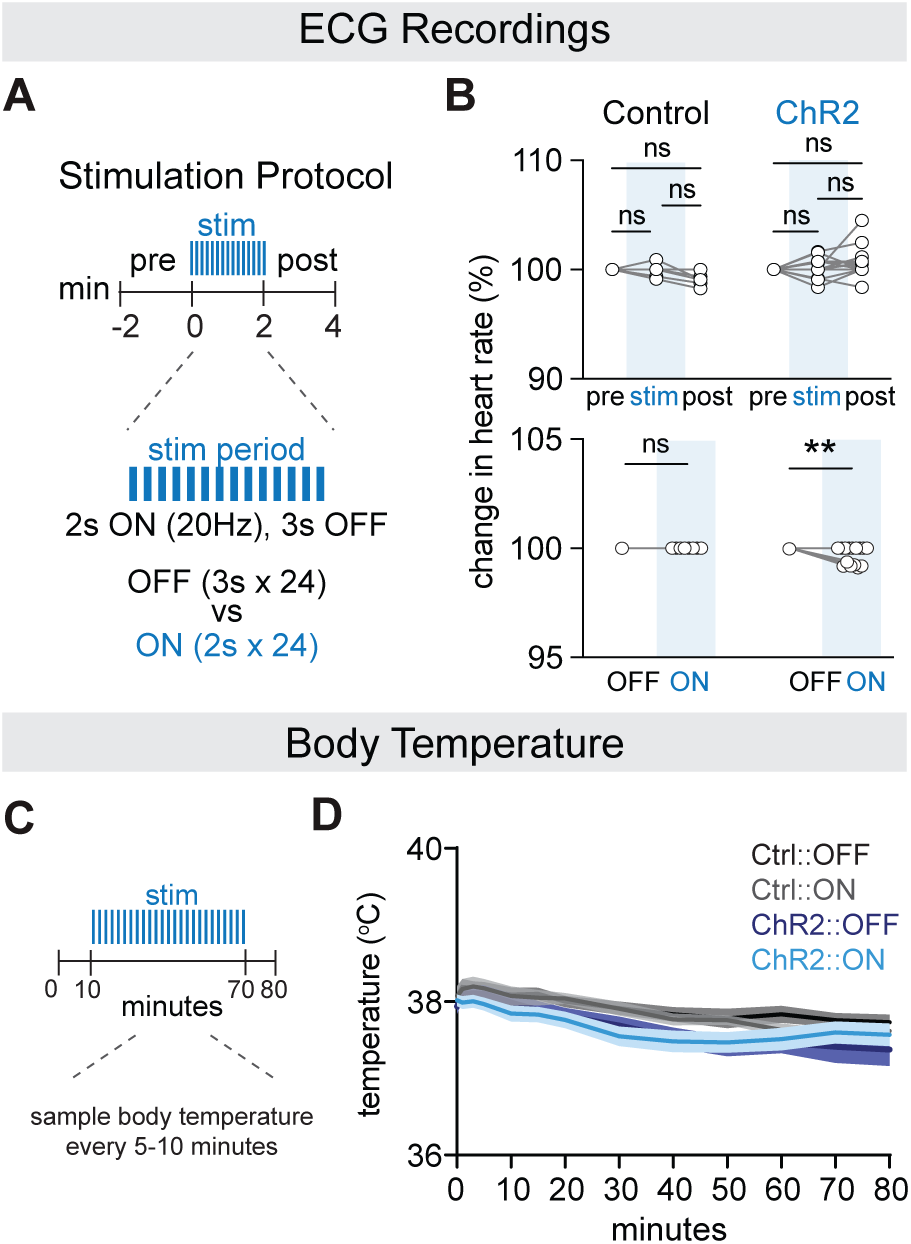
ChR2 stimulation of Npff neurons causes a minimal (∼0.3%) decrease in heart rate and no change in body temperature. **A.** Schematic of the stimulation protocol for heart rate recordings: 2-min pre- and post stimulation periods flanking a 2-min stim period of alternating 2 s ON (20 Hz) / 3 s OFF epochs, repeated 24 times. **B.** Top, no pairwise comparisons between pre-, stim- and post-periods was significant in either control or ChR2 mice. Bottom, quantification of percent change in heart rate during OFF vs. ON epochs within the stim period revealed a small (∼0.3%) decrease in ChR2 mice during ON epochs. No effect was observed in control mice. **C.** Schematic of the stimulation protocol for body temperature recordings. Stimulation began at minute 10 and continued through minute 70, and body temperature was sampled every 5-10 minutes over the 80-minute session. **D.** Body temperature declined minimally and comparably across all four groups over the course of the session, with no evidence of an effect of stimulation in ChR2 mice. **p<0.01, ns, not significant. Data are mean ± s.e.m. Each dot represents a single mouse.

**Supplementary Figure 8.**
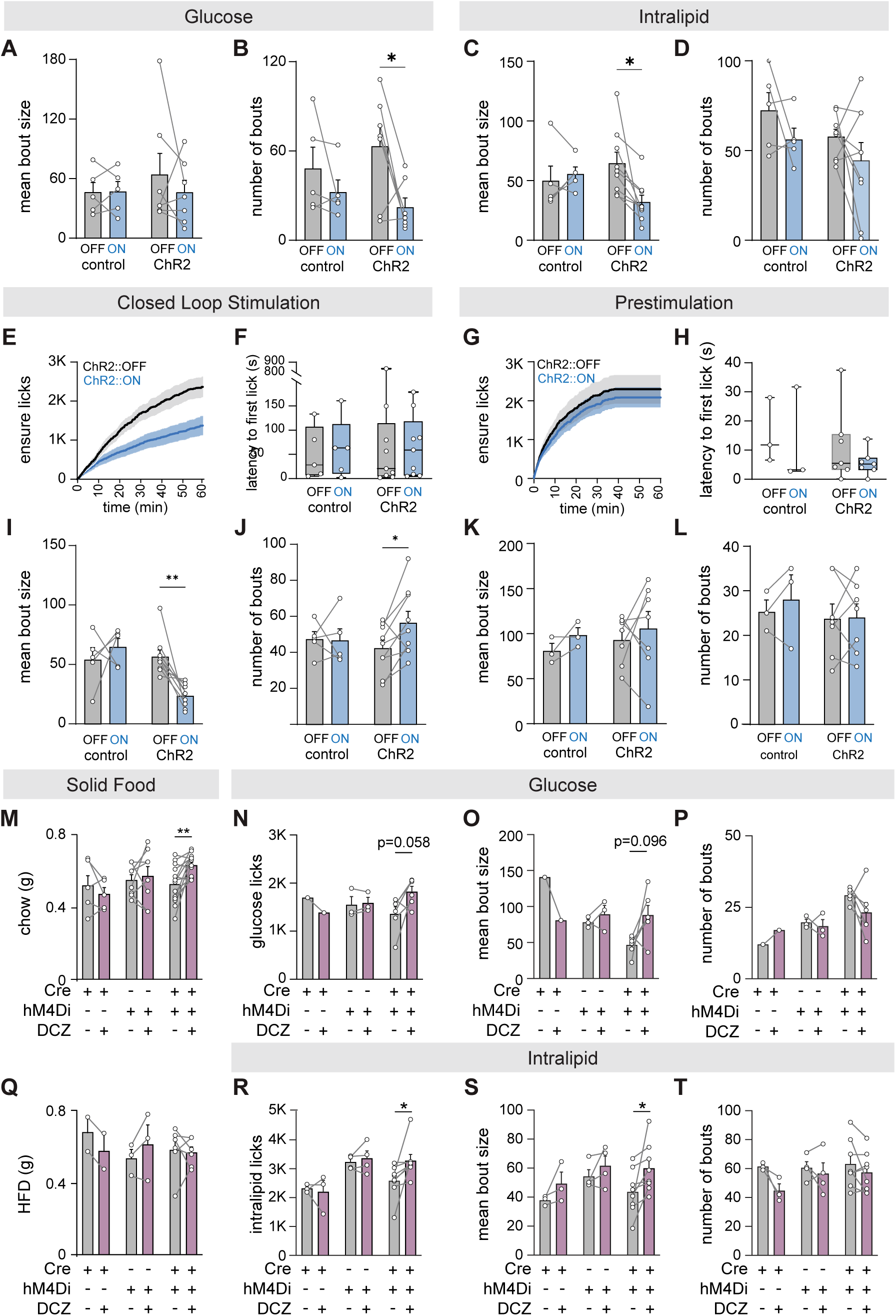
NPFF neuron stimulation and inhibition alter feeding primarily through changes in bout size. **A–D.** Mean bout size (A, C) and number of bouts (B, D) during glucose (A, B) and intralipid (C, D) consumption in control and ChR2 mice during laser OFF and ON conditions. **E, G.** Cumulative Ensure licks over time in ChR2 mice during laser OFF and ON conditions for closed-loop (E) and prestimulation (G) sessions. **F, H.** Latency to first lick in control and ChR2 mice during laser OFF and ON conditions for closed-loop (F) and prestimulation (H) sessions. **I, J.** Mean bout size (I) and number of bouts (J) during closed-loop stimulation in control and ChR2 mice during laser OFF and ON conditions. **K, L.** Mean bout size (K) and number of bouts (L) during prestimulation in control and ChR2 mice during laser OFF and ON conditions. **M, Q.** Chow (M) and HFD (Q) consumption (g) during a 1 h fast-refeed in control and *Npff^Cre^::hM4Di* mice. The control group was further split into Cre-only and hM4Di-only groups. **N–P.** Glucose licks (N), mean bout size (O), and number of bouts (P) in control and *Npff^Cre^::hM4Di* mice following vehicle or DCZ injection. **R–T.** Intralipid licks (R), mean bout size (S), and number of bouts (T) in control and *Npff^Cre^::hM4Di* mice following vehicle or DCZ injection. *p < 0.05, **p < 0.01, ns, not significant. Data are mean ± s.e.m. Each dot represents a single mouse.

## Notes

### Competing Interest Statement

The authors have declared no competing interest.

### Summary of Updates

The supplemental videos were missing from the original submission. They are now included.

## References

1. Müller, M., Canfora, E.E., and Blaak, E.E. (2018). Gastrointestinal Transit Time, Glucose Homeostasis and Metabolic Health: Modulation by Dietary Fibers. Nutrients 10, 275. 10.3390/nu10030275.

2. Huizinga, J.D., and Lammers, W.J.E.P. (2009). Gut peristalsis is governed by a multitude of cooperating mechanisms. Am J Physiol Gastrointest Liver Physiol 296, G1–8. 10.1152/ajpgi.90380.2008.

3. Keller, J., Bassotti, G., Clarke, J., Dinning, P., Fox, M., Grover, M., Hellström, P.M., Ke, M., Layer, P., Malagelada, C., et al. (2018). Advances in the diagnosis and classification of gastric and intestinal motility disorders. Nat Rev Gastroenterol Hepatol 15, 291–308. 10.1038/nrgastro.2018.7.

4. Camilleri, M. (2006). Integrated upper gastrointestinal response to food intake. Gastroenterology 131, 640–658. 10.1053/j.gastro.2006.03.023.

5. Goyal, R.K., Guo, Y., and Mashimo, H. (2019). Advances in the physiology of gastric emptying. Neurogastroenterol Motil 31, e13546. 10.1111/nmo.13546.

6. Marathe, C.S., Rayner, C.K., Jones, K.L., and Horowitz, M. (2013). Relationships Between Gastric Emptying, Postprandial Glycemia, and Incretin Hormones. Diabetes Care 36, 1396–1405. 10.2337/dc12-1609.

7. Camilleri, M., Chedid, V., Ford, A.C., Haruma, K., Horowitz, M., Jones, K.L., Low, P.A., Park, S.-Y., Parkman, H.P., and Stanghellini, V. (2018). Gastroparesis. Nat Rev Dis Primers 4, 41. 10.1038/s41572-018-0038-z.

8. Tack, J., Arts, J., Caenepeel, P., De Wulf, D., and Bisschops, R. (2009). Pathophysiology, diagnosis and management of postoperative dumping syndrome. Nat Rev Gastroenterol Hepatol 6, 583–590. 10.1038/nrgastro.2009.148.

9. Anvari, M., Dent, J., Malbert, C., and Jamieson, G.G. (1995). Mechanics of pulsatile transpyloric flow in the pig. The Journal of Physiology 488, 193–202. 10.1113/jphysiol.1995.sp020957.

10. Heddle, R., Dent, J., Read, N.W., Houghton, L.A., Toouli, J., Horowitz, M., Maddern, G.J., and Downton, J. (1988). Antropyloroduodenal motor responses to intraduodenal lipid infusion in healthy volunteers. Am J Physiol 254, G671–679. 10.1152/ajpgi.1988.254.5.G671.

11. Hou, X., Yin, J., Liu, J., Pasricha, P.J., and Chen, J.D.Z. (2005). In Vivo Gastric and Intestinal Slow Waves in W/Wv Mice. Dig Dis Sci 50, 1335–1341. 10.1007/s10620-005-2783-6.

12. Savoye-Collet, C., Savoye, G., and Smout, A. (2003). Determinants of transpyloric fluid transport: a study using combined real-time ultrasound, manometry, and impedance recording. Am J Physiol Gastrointest Liver Physiol 285, G1147–1152. 10.1152/ajpgi.00208.2003.

13. Code, C.F., and Marlett, J.A. (1975). The interdigestive myo-electric complex of the stomach and small bowel of dogs. The Journal of Physiology 246, 289–309. 10.1113/jphysiol.1975.sp010891.

14. Zsombok, A., and Smith, B.N. (2009). Plasticity of central autonomic neural circuits in diabetes. Biochim Biophys Acta 1792, 423–431. 10.1016/j.bbadis.2008.12.001.

15. Usai-Satta, P., Bellini, M., Morelli, O., Geri, F., Lai, M., and Bassotti, G. (2020). Gastroparesis: New insights into an old disease. World J Gastroenterol 26, 2333–2348. 10.3748/wjg.v26.i19.2333.

16. Szurszewski, J.H. (1998). A 100-year perspective on gastrointestinal motility. Am J Physiol 274, G447–453. 10.1152/ajpgi.1998.274.3.G447.

17. Der-Silaphet, T., Malysz, J., Hagel, S., Larry Arsenault, A., and Huizinga, J.D. (1998). Interstitial cells of cajal direct normal propulsive contractile activity in the mouse small intestine. Gastroenterology 114, 724–736. 10.1016/s0016-5085(98)70586-4.

18. Sanders, K.M., Koh, S.D., Ro, S., and Ward, S.M. (2012). Regulation of gastrointestinal motility—insights from smooth muscle biology. Nat Rev Gastroenterol Hepatol 9, 633–645. 10.1038/nrgastro.2012.168.

19. Spencer, N.J., and Hu, H. (2020). Enteric nervous system: sensory transduction, neural circuits and gastrointestinal motility. Nat Rev Gastroenterol Hepatol 17, 338–351. 10.1038/s41575-020-0271-2.

20. Bayliss, W.M., and Starling, E.H. (1899). The movements and innervation of the small intestine. J Physiol 24, 99–143. 10.1113/jphysiol.1899.sp000752.

21. Furness, J.B. (2012). The enteric nervous system and neurogastroenterology. Nat Rev Gastroenterol Hepatol 9, 286–294. 10.1038/nrgastro.2012.32.

22. Sanders, K.M., Koh, S.D., and Ward, S.M. (2006). Interstitial cells of cajal as pacemakers in the gastrointestinal tract. Annu Rev Physiol 68, 307–343. 10.1146/annurev.physiol.68.040504.094718.

23. Travagli, R.A., Hermann, G.E., Browning, K.N., and Rogers, R.C. (2006). Brainstem Circuits Regulating Gastric Function. Annu Rev Physiol 68, 279–305. 10.1146/annurev.physiol.68.040504.094635.

24. Travagli, R.A., and Anselmi, L. (2016). Vagal neurocircuitry and its influence on gastric motility. Nat Rev Gastroenterol Hepatol 13, 389–401. 10.1038/nrgastro.2016.76.

25. Cannon, W.B., and Lieb, C.W. (1911). The receptive relaxation of the stomach. American Journal of Physiology-Legacy Content 29, 267–273. 10.1152/ajplegacy.1911.29.2.267.

26. De Ponti, F., Azpiroz, F., and Malagelada, J.R. (1987). Reflex gastric relaxation in response to distention of the duodenum. Am J Physiol 252, G595–601. 10.1152/ajpgi.1987.252.5.G595.

27. Reynolds, R.P., El-Sharkawy, T.Y., and Diamant, N.E. (1984). Lower esophageal sphincter function in the cat: role of central innervation assessed by transient vagal blockade. Am J Physiol 246, G666–674. 10.1152/ajpgi.1984.246.6.G666.

28. Rogers, R.C., Hermann, G.E., and Travagli, R.A. (1999). Brainstem pathways responsible for oesophageal control of gastric motility and tone in the rat. J Physiol 514, 369–383. 10.1111/j.1469-7793.1999.369ae.x.

29. Gillis, R.A., Dezfuli, G., Bellusci, L., Vicini, S., and Sahibzada, N. (2021). Brainstem Neuronal Circuitries Controlling Gastric Tonic and Phasic Contractions: A Review. Cell Mol Neurobiol 42, 333–360. 10.1007/s10571-021-01084-5.

30. Browning, K.N., and Travagli, R.A. (2014). Central Nervous System Control of Gastrointestinal Motility and Secretion and Modulation of Gastrointestinal Functions. Comprehensive Physiology 4, 1339–1368. 10.1002/j.2040-4603.2014.tb00587.x.

31. Ran, C., Boettcher, J.C., Kaye, J.A., Gallori, C.E., and Liberles, S.D. (2022). A brainstem map for visceral sensations. Nature 609, 320–326. 10.1038/s41586-022-05139-5.

32. Bai, L., Mesgarzadeh, S., Ramesh, K.S., Huey, E.L., Liu, Y., Gray, L.A., Aitken, T.J., Chen, Y., Beutler, L.R., Ahn, J.S., et al. (2019). Genetic Identification of Vagal Sensory Neurons That Control Feeding. Cell 179, 1129–1143.e23. 10.1016/j.cell.2019.10.031.

33. Williams, E.K., Chang, R.B., Strochlic, D.E., Umans, B.D., Lowell, B.B., and Liberles, S.D. (2016). Sensory Neurons that Detect Stretch and Nutrients in the Digestive System. Cell 166, 209–221. 10.1016/j.cell.2016.05.011.

34. Ludwig, M.Q., Cheng, W., Gordian, D., Lee, J., Paulsen, S.J., Hansen, S.N., Egerod, K.L., Barkholt, P., Rhodes, C.J., Secher, A., et al. (2021). A genetic map of the mouse dorsal vagal complex and its role in obesity. Nat Metab 3, 530–545. 10.1038/s42255-021-00363-1.

35. Camilleri, M., and Linden, D.R. (2016). Measurement of Gastrointestinal and Colonic Motor Functions in Humans and Animals. Cell Mol Gastroenterol Hepatol 2, 412–428. 10.1016/j.jcmgh.2016.04.003.

36. Hatanaka, S., Kondoh, M., Kawarabayashi, K., and Furuhama, K. (1994). The measurement of gastric emptying in conscious rats by monitoring serial changes in serum acetaminophen level. J Pharmacol Toxicol Methods 31, 161–165. 10.1016/1056-8719(94)90079-5.

37. Poli, E., and Pozzoli, C. (2010). Assessment of Gastrointestinal Propulsive Activity Using Three Different Models of Peristalsis In Vivo in the Mouse. Current Protocols in Toxicology 46, 21.9.1–21.9.20. 10.1002/0471140856.tx2109s46.

38. Kendig, D.M., Hurst, N.R., and Grider, J.R. (2016). Spatiotemporal Mapping of Motility in Ex Vivo Preparations of the Intestines. J Vis Exp, 53263. 10.3791/53263.

39. Lentle, R.G., and Hulls, C.M. (2018). Quantifying Patterns of Smooth Muscle Motility in the Gut and Other Organs With New Techniques of Video Spatiotemporal Mapping. Front Physiol 9, 338. 10.3389/fphys.2018.00338.

40. Spencer, N.J., Costa, M., Hibberd, T.J., and Wood, J.D. (2021). Advances in colonic motor complexes in mice. American Journal of Physiology-Gastrointestinal and Liver Physiology 320, G12–G29. 10.1152/ajpgi.00317.2020.

41. Ly, T., Yi, X., Lee, G.R., Grove, J.C., Sibih, Y.E., Oh, J.Y., Qiu, L., Sivakumar, N., and Knight, Z.A. (2025). Cellular coding of ingestion in the caudal brainstem. Preprint at bioRxiv, 10.64898/2025.11.30.691333 10.64898/2025.11.30.691333.

42. Ly, T., Oh, J.Y., Sivakumar, N., Shehata, S., La Santa Medina, N., Huang, H., Liu, Z., Fang, W., Barnes, C., Dundar, N., et al. (2023). Sequential appetite suppression by oral and visceral feedback to the brainstem. Nature 624, 130–137. 10.1038/s41586-023-06758-2.

43. Lever, T.E., Braun, S.M., Brooks, R.T., Harris, R.A., Littrell, L.L., Neff, R.M., Hinkel, C.J., Allen, M.J., and Ulsas, M.A. (2015). Adapting Human Videofluoroscopic Swallow Study Methods to Detect and Characterize Dysphagia in Murine Disease Models. J Vis Exp, 52319. 10.3791/52319.

44. Essner, R.A., Ruda, K., Choh, H.J., Kucukdereli, H., Amsalem, O., Edelhaus, J., Grødem, S., Lensjø, K.K., Lever, T.E., and Andermann, M.L. (2025). Brainstem sensing of multiple body signals during food consumption. Preprint at bioRxiv, 10.1101/2025.04.28.651046 10.1101/2025.04.28.651046.

45. Cannon, W.B. (1898). Movements of the Stomach, Studied by Means of the Röntgen Rays. J Boston Soc Med Sci 2, 59–66.

46. Huizinga, J.D., and Chen, J.-H. (2014). The myogenic and neurogenic components of the rhythmic segmentation motor patterns of the intestine. Front. Neurosci. 8. 10.3389/fnins.2014.00078.

47. Berthoud, H.R., and Powley, T.L. (1992). Vagal afferent innervation of the rat fundic stomach: morphological characterization of the gastric tension receptor. J Comp Neurol 319, 261–276. 10.1002/cne.903190206.

48. Altschuler, S.M., Bao, X., Bieger, D., Hopkins, D.A., and Miselis, R.R. (1989). Viscerotopic representation of the upper alimentary tract in the rat: Sensory ganglia and nuclei of the solitary and spinal trigeminal tracts. Journal of Comparative Neurology 283, 248–268. 10.1002/cne.902830207.

49. Travagli, R.A., Hermann, G.E., Browning, K.N., and Rogers, R.C. (2003). Musings on the wanderer: what’s new in our understanding of vago-vagal reflexes? III. Activity-dependent plasticity in vago-vagal reflexes controlling the stomach. Am J Physiol Gastrointest Liver Physiol 284, G180–187. 10.1152/ajpgi.00413.2002.

50. Yao, Z., van Velthoven, C.T.J., Kunst, M., Zhang, M., McMillen, D., Lee, C., Jung, W., Goldy, J., Abdelhak, A., Aitken, M., et al. (2023). A high-resolution transcriptomic and spatial atlas of cell types in the whole mouse brain. Nature 624, 317–332. 10.1038/s41586-023-06812-z.

51. López-Cruz, A., Burgos, N.S.F., Hakimi, A.M., Xie, K., Mao, M., Kaur, M., Spina, A., Choi, K., Qiu, L., Lever, T.E., et al. (2026). Functional segregation of body-brain signals in the area postrema. Preprint at bioRxiv, 10.64898/2026.06.15.732473 10.64898/2026.06.15.732473.

52. Gasparini, S., Zhu, H., Munuzuri, A.S.P., Leuang, M.L., Fazan, F.S., and Geerling, J.C. (2026). Molecular Ontology Predicts Output Connections From the Nucleus of the Solitary Tract. J Comp Neurol 534, e70188. 10.1002/cne.70188.

53. Callegari, E., Malhotra, B., Bungay, P.J., Webster, R., Fenner, K.S., Kempshall, S., LaPerle, J.L., Michel, M.C., and Kay, G.G. (2011). A comprehensive non-clinical evaluation of the CNS penetration potential of antimuscarinic agents for the treatment of overactive bladder. British Journal of Clinical Pharmacology 72, 235–246. 10.1111/j.1365-2125.2011.03961.x.

54. Yoshida, A., Maruyama, S., Fukumoto, D., Tsukada, H., Ito, Y., and Yamada, S. (2010). Noninvasive evaluation of brain muscarinic receptor occupancy of oxybutynin, darifenacin and imidafenacin in rats by positron emission tomography. Life Sciences 87, 175–180. 10.1016/j.lfs.2010.06.008.

55. Matsui, M., Motomura, D., Fujikawa, T., Jiang, J., Takahashi, S., Manabe, T., and Taketo, M.M. (2002). Mice lacking M2 and M3 muscarinic acetylcholine receptors are devoid of cholinergic smooth muscle contractions but still viable. J Neurosci 22, 10627–10632. 10.1523/JNEUROSCI.22-24-10627.2002.

56. Tanahashi, Y., Komori, S., Matsuyama, H., Kitazawa, T., and Unno, T. (2021). Functions of Muscarinic Receptor Subtypes in Gastrointestinal Smooth Muscle: A Review of Studies with Receptor-Knockout Mice. Int J Mol Sci 22, 926. 10.3390/ijms22020926.

57. Unno, T., Matsuyama, H., Izumi, Y., Yamada, M., Wess, J., and Komori, S. (2006). Roles of M2 and M3 muscarinic receptors in cholinergic nerve-induced contractions in mouse ileum studied with receptor knockout mice. Br J Pharmacol 149, 1022–1030. 10.1038/sj.bjp.0706955.

58. Bharucha, A.E., Ravi, K., and Zinsmeister, A.R. (2010). Comparison of selective M3 and nonselective muscarinic receptor antagonists on gastrointestinal transit and bowel habits in humans. American Journal of Physiology-Gastrointestinal and Liver Physiology 299, G215–G219. 10.1152/ajpgi.00072.2010.

59. Eglen, R.M., Choppin, A., and Watson, N. (2001). Therapeutic opportunities from muscarinic receptor research. Trends in Pharmacological Sciences 22, 409–414. 10.1016/S0165-6147(00)01737-5.

60. Takeuchi, T., Fujinami, K., Goto, H., Fujita, A., Taketo, M.M., Manabe, T., Matsui, M., and Hata, F. (2005). Roles of M2 and M4 Muscarinic Receptors in Regulating Acetylcholine Release From Myenteric Neurons of Mouse Ileum. Journal of Neurophysiology 93, 2841–2848. 10.1152/jn.00986.2004.

61. Kilbinger, H., Halim, S., Lambrecht, G., Weiler, W., and Wessler, I. (1984). Comparison of affinities of muscarinic antagonists to pre- and postjunctional receptors in the guinea-pig ileum. European Journal of Pharmacology 103, 313–320. 10.1016/0014-2999(84)90492-8.

62. Salvador, A.F.M., Golynker, I., Betley, J.N., and Thaiss, C.A. (2026). Intestinal interoception: A nexus of environment-body-brain interactions. Neuron 114, 583–600. 10.1016/j.neuron.2025.11.026.

63. Cannon, W.B. (1909). The Influence of Emotional States on the Functions of the Alimentary Canal.

64. Furness, J.B., Callaghan, B.P., Rivera, L.R., and Cho, H.-J. (2014). The enteric nervous system and gastrointestinal innervation: integrated local and central control. Adv Exp Med Biol 817, 39–71. 10.1007/978-1-4939-0897-4_3.

65. Shahidullah, M., Kennedy, T.L., and Parks, T.G. (1975). The vagus, the duodenal brake, and gastric emptying. Gut 16, 331–336. 10.1136/gut.16.5.331.

66. Treacy, P.J., Jamieson, G.G., Dent, J., Devitt, P.G., and Heddle, R. (1992). Duodenal intramural nerves in control of pyloric motility and gastric emptying. Am J Physiol 263, G1–5. 10.1152/ajpgi.1992.263.1.G1.

67. Treacy, P.J., Jamieson, G.G., and Dent, J. (1996). The Effect of Duodenal Distension Upon Antro-Pyloric Motility and Liquid Gastric Emptying in Pigs. Australian and New Zealand Journal of Surgery 66, 37–40. 10.1111/j.1445-2197.1996.tb00698.x.

68. Spiller, R.C., Trotman, I.F., Higgins, B.E., Ghatei, M.A., Grimble, G.K., Lee, Y.C., Bloom, S.R., Misiewicz, J.J., and Silk, D.B. (1984). The ileal brake--inhibition of jejunal motility after ileal fat perfusion in man. Gut 25, 365–374. 10.1136/gut.25.4.365.

69. Zhang, T., Perkins, M.H., Chang, H., Han, W., and Araujo, I.E. de (2022). An inter-organ neural circuit for appetite suppression. Cell 185, 2478–2494.e28. 10.1016/j.cell.2022.05.007.

70. Maljaars, P.W.J., Peters, H.P.F., Mela, D.J., and Masclee, A. a. M. (2008). Ileal brake: a sensible food target for appetite control. A review. Physiol Behav 95, 271–281. 10.1016/j.physbeh.2008.07.018.

71. Read, N.W., McFarlane, A., Kinsman, R.I., Bates, T.E., Blackhall, N.W., Farrar, G.B., Hall, J.C., Moss, G., Morris, A.P., and O’Neill, B. (1984). Effect of infusion of nutrient solutions into the ileum on gastrointestinal transit and plasma levels of neurotensin and enteroglucagon. Gastroenterology 86, 274–280.

72. Sabatini, P.V., Frikke-Schmidt, H., Arthurs, J., Gordian, D., Patel, A., Rupp, A.C., Adams, J.M., Wang, J., Beck Jørgensen, S., Olson, D.P., et al. (2021). GFRAL-expressing neurons suppress food intake via aversive pathways. Proceedings of the National Academy of Sciences 118, e2021357118. 10.1073/pnas.2021357118.

73. Yacawych, W.T., Wang, Y., Zhou, G., Hassan, S., Kernodle, S., Sass, F., DeVaux, M., Wu, I., Rupp, A., Tomlinson, A.J., et al. (2025). A single dorsal vagal complex circuit mediates the aversive and anorectic responses to GLP1R agonists. Preprint at bioRxiv, 10.1101/2025.01.21.634167 10.1101/2025.01.21.634167.

74. Lewin, A.E., Vicini, S., Richardson, J., Dretchen, K.L., Gillis, R.A., and Sahibzada, N. (2016). Optogenetic and pharmacological evidence that somatostatin-GABA neurons are important regulators of parasympathetic outflow to the stomach. The Journal of Physiology 594, 2661–2679. 10.1113/JP272069.

75. Bellusci, L., DuBar, S.N.G., Kuah, M., Castellano, D., Muralidaran, V., Jones, E., Rozeboom, A.M., Gillis, R.A., Vicini, S., and Sahibzada, N. (2022). Interactions between Brainstem Neurons That Regulate the Motility to the Stomach. J. Neurosci. 42, 5212–5228. 10.1523/JNEUROSCI.0419-22.2022.

76. Cruz, M.T., Murphy, E.C., Sahibzada, N., Verbalis, J.G., and Gillis, R.A. (2007). A reevaluation of the effects of stimulation of the dorsal motor nucleus of the vagus on gastric motility in the rat. American Journal of Physiology-Regulatory, Integrative and Comparative Physiology 292, R291–R307. 10.1152/ajpregu.00863.2005.

77. Taché, Y., Raybould, H., and Wei, J.Y. (1991). Central and peripheral actions of calcitonin gene-related peptide on gastric secretory and motor function. Adv Exp Med Biol 298, 183–198. 10.1007/978-1-4899-0744-8_17.

78. Stengel, A., and Taché, Y. (2010). Corticotropin-releasing factor signaling and visceral response to stress. Exp Biol Med (Maywood) 235, 1168–1178. 10.1258/ebm.2010.009347.

79. Ludwig, M.Q., Coester, B., Gordian, D., Hassan, S., Tomlinson, A.J., Toure, M.H., Christensen, O.P., Lommi, G., Moltke-Prehn, A., Brown, J.M., et al. (2026). A cross-species atlas of the dorsal vagal complex reveals neural mediators of the effects of cagrilintide on energy balance. Nat Metab 8, 1350–1367. 10.1038/s42255-026-01539-3.

80. Ruud, L.E., Font-Gironès, F., Zajdel, J., Kern, L., Teixidor-Deulofeu, J., Mannerås-Holm, L., Carreras, A., Becattini, B., Björefeldt, A., Hanse, E., et al. (2024). Activation of GFRAL+ neurons induces hypothermia and glucoregulatory responses associated with nausea and torpor. Cell Reports 43. 10.1016/j.celrep.2024.113960.

81. Zhang, X., Fogel, R., and Renehan, W.E. (1995). Relationships between the morphology and function of gastric- and intestine-sensitive neurons in the nucleus of the solitary tract. Journal of Comparative Neurology 363, 37–52. 10.1002/cne.903630105.

82. Gasparini, S., Almeida-Pereira, G., Munuzuri, A.S.P., Resch, J.M., and Geerling, J.C. (2024). Molecular Ontology of the Nucleus of Solitary Tract. Journal of Comparative Neurology 532, e70004. 10.1002/cne.70004.

83. Kivipelto, L., Aarnisalo, A., and Panula, P. (1992). Neuropeptide FF is colocalized with catecholamine-synthesizing enzymes in neurons of the nucleus of the solitary tract. Neuroscience Letters 143, 190–194. 10.1016/0304-3940(92)90263-7.

84. Browning, K.N., Travagli, R.A., and Pellegrini, C. (2026). Central control of gastrointestinal functions in health and disease. Physiological Reviews 106, 971–1020. 10.1152/physrev.00010.2025.

85. Wang, X., Alkaabi, F., Choi, M., Di Natale, M.R., Scheven, U.M., Noll, D.C., Furness, J.B., and Liu, Z. (2024). Surface mapping of gastric motor functions using MRI: a comparative study between humans and rats. American Journal of Physiology-Gastrointestinal and Liver Physiology 327, G345–G359. 10.1152/ajpgi.00045.2024.

86. Hennig, G.W., and Spencer, N.J. (2018). Chapter 21 - Physiology of Gastric Motility Patterns. In Physiology of the Gastrointestinal Tract (Sixth Edition), H. M. Said, ed. (Academic Press), pp. 469–484. 10.1016/B978-0-12-809954-4.00021-9.

87. Richardson, J., Dezfuli, G., Mangel, A.W., Gillis, R.A., Vicini, S., and Sahibzada, N. (2023). CNS sites controlling the gastric pyloric sphincter: Neuroanatomical and functional study in the rat. Journal of Comparative Neurology 531, 1562–1581. 10.1002/cne.25530.

88. Chang, H.Y., Mashimo, H., and Goyal, R.K. (2003). IV. Current concepts of vagal efferent projections to the gut. American Journal of Physiology-Gastrointestinal and Liver Physiology 284, G357–G366. 10.1152/ajpgi.00478.2002.

89. Malbert, C.H., Mathis, C., and Laplace, J.P. (1994). Vagal control of transpyloric flow and pyloric resistance. Digest Dis Sci 39, 24S–27S. 10.1007/BF02300364.

90. Pal, A., Indireshkumar, K., Schwizer, W., Abrahamsson, B., Fried, M., and Brasseur, J.G. (2004). Gastric flow and mixing studied using computer simulation. Proc Biol Sci 271, 2587–2594. 10.1098/rspb.2004.2886.

91. Wang, Y., Chen, F., Shi, H., Jiang, J., Li, H., Qin, B., and Li, Y. (2015). Extrinsic ghrelin in the paraventricular nucleus increases small intestinal motility in rats by activating central growth hormone secretagogue and enteric cholinergic receptors. Peptides 74, 43–49. 10.1016/j.peptides.2015.09.009.

92. Martínez, V., Wang, L., Rivier, J., Grigoriadis, D., and Taché, Y. (2004). Central CRF, urocortins and stress increase colonic transit via CRF1 receptors while activation of CRF2 receptors delays gastric transit in mice. J Physiol 556, 221–234. 10.1113/jphysiol.2003.059659.

93. Taché, Y., Garrick, T., and Raybould, H. (1990). Central nervous system action of peptides to influence gastrointestinal motor function. Gastroenterology 98, 517–528. 10.1016/0016-5085(90)90849-v.

94. Berthoud, H.R., Carlson, N.R., and Powley, T.L. (1991). Topography of efferent vagal innervation of the rat gastrointestinal tract. American Journal of Physiology-Regulatory, Integrative and Comparative Physiology 260, R200–R207. 10.1152/ajpregu.1991.260.1.R200.

95. Wang, X., Alkaabi, F., Cornett, A., Choi, M., Scheven, U.M., Di Natale, M.R., Furness, J.B., and Liu, Z. (2026). Magnetic Resonance Imaging of Gastric Motility in Conscious Rats. Neurogastroenterology & Motility 38, e14982. 10.1111/nmo.14982.

96. Ravi, N., Gabeur, V., Hu, Y.-T., Hu, R., Ryali, C., Ma, T., Khedr, H., Rädle, R., Rolland, C., Gustafson, L., et al. (2024). SAM 2: Segment Anything in Images and Videos. Preprint at arXiv, 10.48550/arXiv.2408.00714 10.48550/arXiv.2408.00714.

